# A systematic evaluation and benchmarking of text summarization methods for biomedical literature: From word-frequency methods to language models

**DOI:** 10.64898/2026.01.09.697335

**Authors:** Fabio Baumgärtel, Enrico Bono, Lucas Fillinger, Louiza Galou, Kinga Keska-Izworska, Samuel M Walter, Peter Andorfer, Klaus Kratochwill, Paul Perco, Matthias Ley

## Abstract

The rapid expansion of biomedical literature demands automated summarization tools that reliably condense research articles into concise, accurate summaries. We benchmarked 62 summarization methods, ranging from frequency-based and TextRank extractors to encoder-decoder models (EDMs) and large language models (LLMs), on 1,000 biomedical abstracts from 20 journals across *ScienceDirect* and *Cell Press*, using author-written highlights as reference summaries. Models were evaluated with a composite suite of lexical, semantic, and factual metrics, including ROUGE, BLEU, METEOR, embedding-based similarity, and factuality scores. General-purpose models (e.g., Mistral, GPT, Llama) achieved the highest overall performance across lexical and semantic dimensions, outperforming reasoning-oriented (e.g., DeepSeek, Magistral) and domain-specific (e.g., BioGPT, BioMistral) models. Notably, medium-sized models outperformed large-scale models, suggesting an optimal balance between model capacity and efficiency, while classical extractive methods lagged behind neural approaches. These findings provide a systematic reference for selecting biomedical summarization tools and highlight that broad pretraining outperforms narrow domain adaptation.

## Introduction

The exponential growth of scientific literature has created a demand for text summarization methods to support scientists in prioritizing articles, extracting relevant information, and interpreting research findings. Automatic text summarization (ATS) methods have evolved from statistical approaches to deep learning-based models, thus becoming increasingly sophisticated and reliable in capturing the essential parts of complex research articles. Although ATS methods have been previously evaluated and described^1,2^, only a few have focused on scientific literature^3,4^. Recent work has begun to benchmark large language models (LLMs) on biomedical summarization, though only for a limited set of models and typically against abstract-based references^5–7^. A systematic comparison from word-frequency methods to state-of-the-art LLMs is still lacking.

Over time, these methods have developed into distinct approaches that differ in how they identify or generate summary content. Early pre-neural summarization was mainly characterized by unsupervised extractive approaches that rank sentences by word or concept frequencies to identify relevant sentences. The field was initially shaped by Luhn’s assumption that recurrent words signal importance^8^ and Edmundson’s use of cue words, title words, and sentence position^9^. Later work adopting weighting term frequency-inverse document frequency (TF-IDF) refined this idea by down-weighting frequent, low-specificity terms^10^, laying the foundation for more advanced methods in scientific text summarization^11^. Graph-based methods judged the importance against the document’s global structure rather than local word counts. TextRank, widely used in the biomedical domain, applies the PageRank algorithm to a sentence-similarity graph to select top-ranked sentences^12^.

The advent of sequence-to-sequence (Seq2seq) frameworks shifted the field toward neural approaches capable of paraphrasing and condensing text using encoder-decoder architectures, originally implemented with recurrent neural networks (RNNs), long short-term memory (LSTM) networks, and gated recurrent units (GRUs)^13,14^, while self-attention enabled parallel processing of sequences and modeling of long-range context^15^. This laid the foundation for transformer architectures that quickly gained popularity in performing a wide range of natural language processing (NLP) tasks. Building on bidirectional encoder representations from transformers (BERT)^16^, several abstractive summarization models have emerged, including bidirectional and auto-regressive transformer (BART), a denoising autoencoder for pretraining Seq2seq models^17^ and its biomedical adaptations^18,19^; the unified text-to-text transfer transformer (T5) model^20^; Pre-training with extracted gap-sentences for abstractive summarization sequence-to-sequence (PEGASUS)^21^, including domain-specific PubMed variants; and Longformer-based models^22^ that can handle longer inputs^23^. These models are typically pretrained independently, while Longformer-based variants may additionally leverage RoBERTa, an optimized version of BERT trained on a larger corpus.

More recently, the field of ATS has quickly moved toward decoder-only architectures at the core of LLMs, which capture semantic relations with greater flexibility and specificity. LLMs can be classified as (i) general-purpose models, which apply broad domain knowledge across diverse NLP tasks, such as the proprietary generative pre-trained transformer (GPT)^24^ and Claude^25^ series and the open-source large language model Meta AI (Llama)^26^ and Gemma^27^ families; (ii) reasoning-oriented models, characterized by logical text understanding through iterative chain-of-thought processing and instruction tuning^28^ such as DeepSeek-R1^29^, Qwen3^30^, and Mistral’s Magistral^31^, together with the reasoning tiers of the proprietary families (e.g., GPT-5 and Claude Opus 4); and (iii) domain-specific models, tailored for specialized tasks or specific scientific domains, such as the biomedical models OpenBioLLM-Llama3^32^, BioGPT^33^, and BioMistral^34^, and SciLitLLM^35^ for scientific literature. The complete set of models evaluated in this study is listed in Table 2 in the Materials and Methods.

Despite this range of approaches, these method families have not been evaluated together on biomedical literature, leaving little guidance on which to choose. This study addresses this gap by systematically evaluating 62 text summarization models, ranging from word-frequency methods to state-of-the-art LLMs, using a dataset of 1,000 biomedical abstracts and corresponding highlights sections as reference summaries. We provide recommendations for selecting appropriate tools to accelerate knowledge discovery in biomedical sciences and discuss the strengths and limitations of each approach.

## Materials and Methods

### Study Design

This study was designed as a systematic evaluation and comparative benchmark of ATS methods for biomedical literature and is reported in accordance with the transparent reporting of a multivariable prediction model for individual prognosis or diagnosis (TRIPOD)-LLM reporting guideline^36^. Summarization quality was assessed quantitatively using a suite of lexical, semantic, and factual metrics, normalized and aggregated into an Overall Performance Score for ranking, and complemented by a blinded expert assessment of a subset of models. Differences across model categories and families were then compared statistically. The evaluation metrics and statistical tests were chosen during the analysis rather than pre-specified in a study protocol.

Figure 1 provides an overview of the complete benchmarking workflow of this study from dataset construction and model inference to metric computation, score aggregation, and statistical analysis.

**Figure 1:**
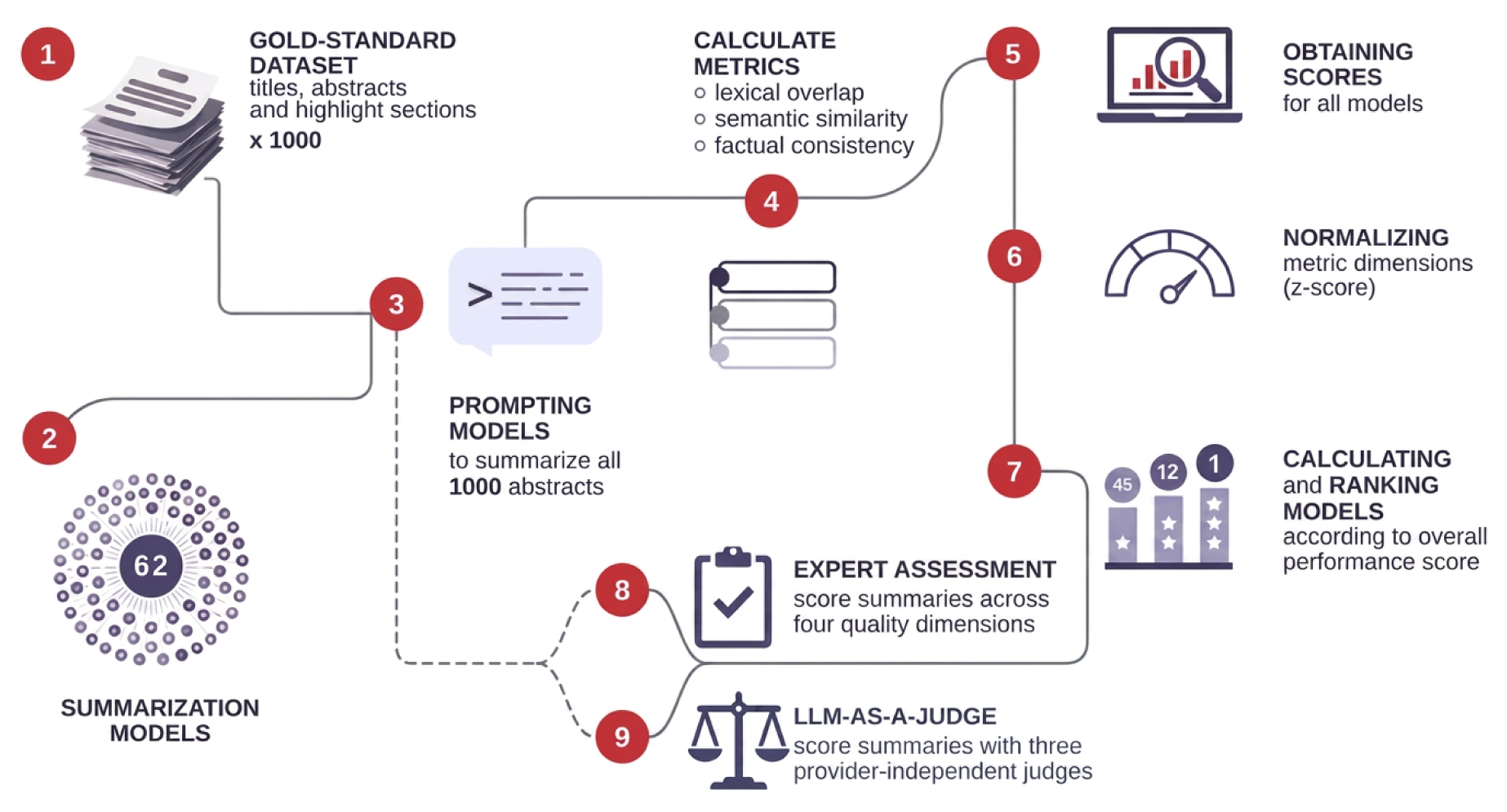
Overview of the benchmarking workflow: (1) Construction of a gold-standard dataset comprising 1,000 biomedical titles, abstracts, and author-provided highlight sections. (2) Suite of 62 diverse summarization models. (3) Uniform prompting of all models to generate summaries for each abstract. (4–5) Computation of summary quality metrics capturing lexical overlap, semantic similarity, and factual consistency. (6) Standardization of metric scores using z-score normalization to ensure comparability across evaluation dimensions. (7) Ranking of models based on overall performance scores. (8) Quality assessment of generated summaries across four quality dimensions (coherence, fluency, relevance, consistency), combining expert human evaluation on a subset of models with (9) LLM-as-a-judge evaluation from three provider-independent judges run across all models.

### Gold-Standard Dataset

We generated a gold-standard benchmarking dataset comprising 1,000 biomedical peer-reviewed articles from *ScienceDirect* and *Cell Press*, as these publishers provide standardized highlights sections - concise bullet points capturing an article’s main findings^37,38^. The concatenated highlights served as reference summaries in our evaluation, while the corresponding titles and abstracts were used as input texts for the summarization task. While summarization is inherently subjective, author-generated highlights represent the most credible and standardized source, ensuring the captured content reflects the study’s intended key messages.

Articles were collected systematically across a variety of journals from the two publishers to ensure coverage of different fields within molecular sciences, including among others drug discovery, genomics, proteomics, biotechnology, and biochemistry. We considered only English-language, research-focused content that provided an author-written highlights section together with a DOI, title, and abstract. From each of the 20 journals, we then selected the 50 most recent eligible articles available at the time of dataset construction, yielding 1,000 papers (Table 1) with publication dates ranging from October 2015 to November 2025. Because selection was performed per journal, journals with fewer highlights-bearing articles contributed some older publications, though 95% of the corpus was published from 2024 onward. Selecting the most recent articles also minimizes overlap with the models’ pretraining data and thus the risk of data leakage.

**Table 1:** Overview of journals included in the gold-standard dataset: Each journal contributed 50 articles, resulting in 500 articles from *ScienceDirect* and 500 articles from *Cell Press*.

| Publisher | Journals |
| --- | --- |
| *ScienceDirect* | Cytokine; Developmental Cell; Drug Discovery Today; FEBS Letters; Gene; Genomics; Journal of Biotechnology; Journal of Molecular Biology; Journal of Proteomics; The International Journal of Biochemistry & Cell Biology |
| *Cell Press* | Cancer Cell; Cell; Cell Chemical Biology; Cell Genomics; Cell Host & Microbe; Cell Metabolism; Cell Reports; Cell Reports Medicine; Cell Stem Cell; Cell Systems |

This setup provided standardized pairs of abstracts and reference summaries subsequently used in our systematic evaluation and benchmarking of ATS methods.

### Summarization Methods

We evaluated 62 models for text summarization, ranging from simple frequency-based algorithms to small language models (SLMs) and LLMs. These models were classified into five categories, as listed in Table 2. We obtained the pre-trained encoder-decoder models (EDMs) through the Hugging Face library, selecting architectures widely used for abstractive summarization to ensure established neural approaches are covered in our benchmark. Additionally, we evaluated a range of SLMs and LLMs, defining SLMs as models with fewer than 10 billion parameters^39^. Both SLMs and LLMs with advanced reasoning capabilities were categorized as “reasoning-oriented” to investigate how multi-step problem solving affects summarization performance. Similarly, models fine-tuned on scientific/biomedical data or specifically tailored for text summarization were classified as “domain-specific” to assess the impact of domain adaptation. Overall, this selection covers various model sizes and release dates, ensuring a representative mix of both, widely adopted and recent, architectures.

**Table 2:**
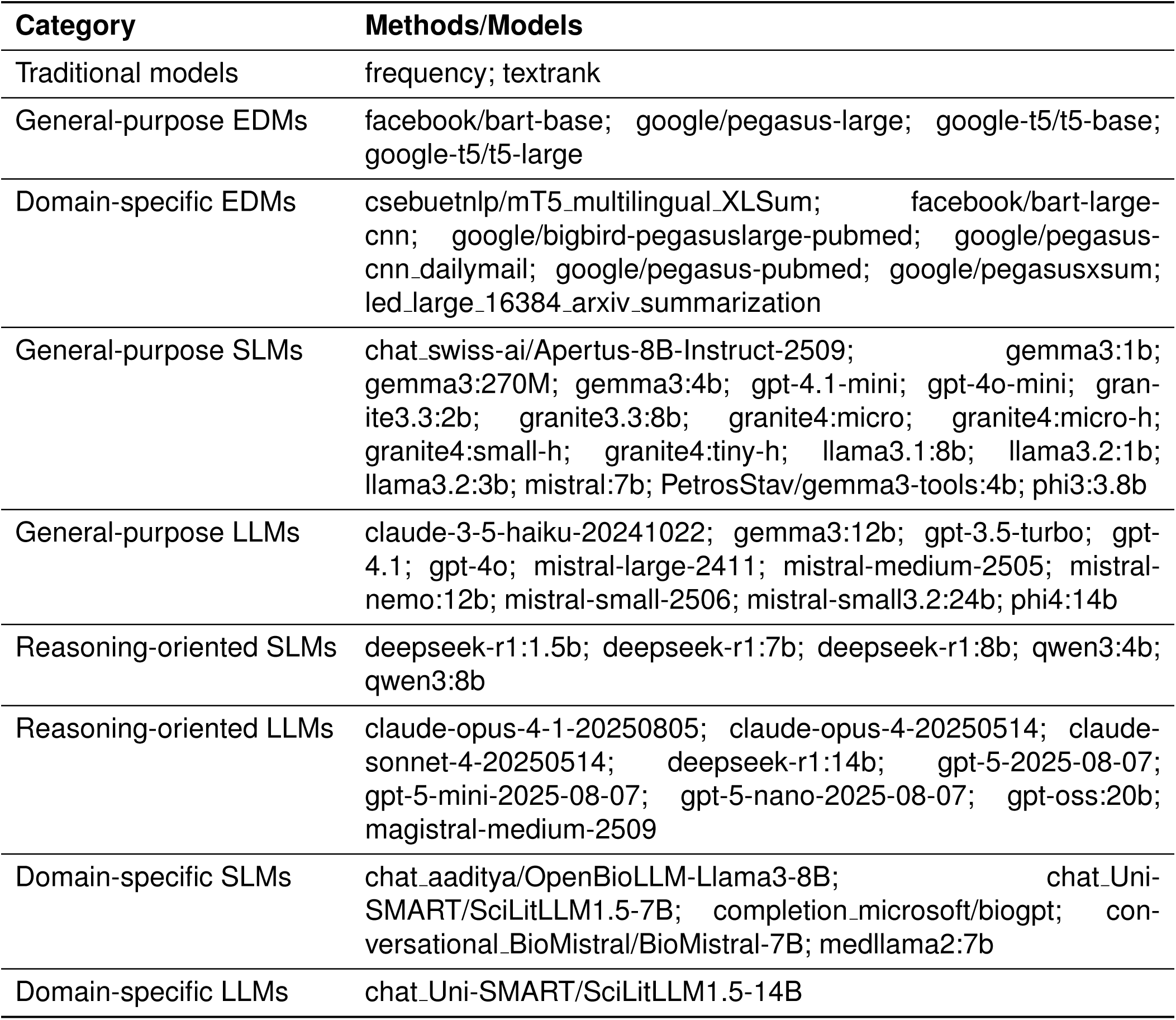
Overview of summarization methods evaluated in this study: Models organized by category.

Exceptionally large models, such as LLaMA 3.1 405B, were excluded to keep the benchmark reproducible on single-GPU setups typical of biomedical research groups. Each of the 62 models was tasked with generating summaries for the 1,000 abstracts in the dataset, resulting in a total of 62,000 generated summaries for evaluation.

### Prompt Design

We used an identical prompt across all prompt-based models to ensure compatibility, asking for concise summaries focused on the main findings of each publication while excluding unnecessary background or methodological details. The prompt was developed iteratively on a small number of sample abstracts and then frozen. Each model received the publication title and abstract as an input and was asked to produce an output of 15–100 words. A post-hoc check confirmed that models adhered well to this length constraint, with the large majority of summaries falling within the target range across methods. A full per-method breakdown is provided in Supplementary Figure 7. If the abstract did not contain any substantive results or conclusions, the model was instructed to return the predefined token INSUFFICIENT FINDINGS. This gave models an explicit exit path for cases where an abstract contained no actual findings to summarize, preventing them from fabricating content simply to fill the response.

The exact prompt used for all models was as follows:

Summarize the provided publication (title and abstract) in 15-100 words.

Key requirements:

- Identify main findings, results, or contributions
- Preserve essential context and nuance
- Exclude background, methods unless crucial to conclusions
- Write concisely and objectively
- Avoid repetition and unnecessary qualifiers

If no substantial findings exist, respond: ’INSUFFICIENT_FINDINGS’

### Evaluation Metrics

As there is no single metric that can fully reflect summary quality, especially in the biomedical field where both coverage of key information and factual correctness are critical, we employed both surface-level metrics based on lexical overlap, referred to as lexical-based metrics, and embedding-level metrics. The latter includes metrics based on semantic similarity, denoted as semantic-based metrics, as well as metrics that evaluate factual consistency.

#### Surface-level Metrics

Surface-level metrics compare the generated summaries with the reference summaries mainly at the word or phrase level. While they do not capture meaning beyond surface overlap, they remain common metrics in summarization research and provide a straightforward foundation for evaluation. We used three recall-oriented understudy for gisting evaluation (ROUGE) variants (ROUGE-1, ROUGE-2, ROUGE-L)^40^, bilingual evaluation understudy (BLEU)^41^, and metric for evaluation of translation with explicit ordering (METEOR)^42^. ROUGE-1 and ROUGE-2 measure how many unigrams (single words) or bigrams (word pairs) from the reference appear in the generated output, while ROUGE-L identifies the longest sequence of words shared between the two. BLEU calculates how many n-grams in the output also occur in the reference, emphasizing precision over recall and applying a brevity penalty to counteract the tendency toward overly short summaries. METEOR extends n-gram matching by considering word stems and synonyms, making it more robust to wording variations. Together, these metrics offer a simple but transparent point of reference.

#### Embedding-based Metrics

To capture similarity beyond surface-level word overlap, we included a set of embedding-based metrics built on pre-trained transformer models. These methods generate vector representations of text, allowing them to capture semantic similarity rather than just word overlap. We measured semantic similarity using BERT-score^43^, which compares two texts by matching their contextual token embeddings and averaging the resulting cosine similarities. Because BERT-score needs a pre-trained encoder to produce these embeddings, we ran it with two well-established choices: Robustly optimized BERT approach (RoBERTa)^44^ and decoding-enhanced BERT with disentangled attention (DeBERTa)^45^. This lets the metrics recognize when two summaries express the same content even if the wording differs.

We further included all-mpnet-base-v2^46^, a transformer model fine-tuned for sentence similarity. Unlike BERT-score, which matches summaries token by token, masked and permuted pre-training (MPNet) was trained with a focus on alignment at the sentence-level. This characteristic makes it a useful complement to the other metrics, as it is particularly sensitive to whether the overall meaning of a reference summary is preserved in the system output.

Finally, to evaluate factual consistency, we included metrics designed to assess whether statements in a generated summary are supported by the source text. These included AlignScore^47^, SummaC^48^, and two variants of MiniCheck (FT5 and 7B)^49^. In contrast to the other metrics, these approaches compare the generated output directly to the source text itself (i.e., the publication abstract) rather than the reference summary (i.e., the highlights section), as factual accuracy can only be assessed relative to the original input text. This addition ensures that our evaluation captures hallucinations and unsupported statements that might otherwise be overlooked.

#### Overall Performance Metric

To comprehensively assess the performance of each model on the summarization task, we employed a multi-metric framework covering three dimensions: lexical (n=5), semantic (n=3), and factual (n=4). To prevent the impact of dimension imbalance, we computed an average score for each dimension as follows:

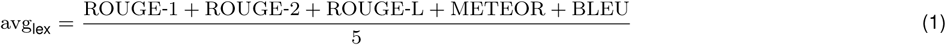

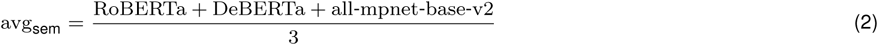

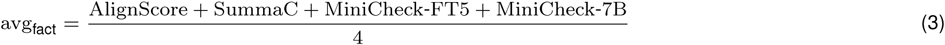

To examine how the different evaluation metrics relate to each other, we computed pairwise Spearman correlation coefficients across all models (Figure 2). Averaging metrics within each dimension ensured that highly correlated measures such as ROUGE-1, ROUGE-2, and ROUGEL do not dominate the overall evaluation.

**Figure 2:**
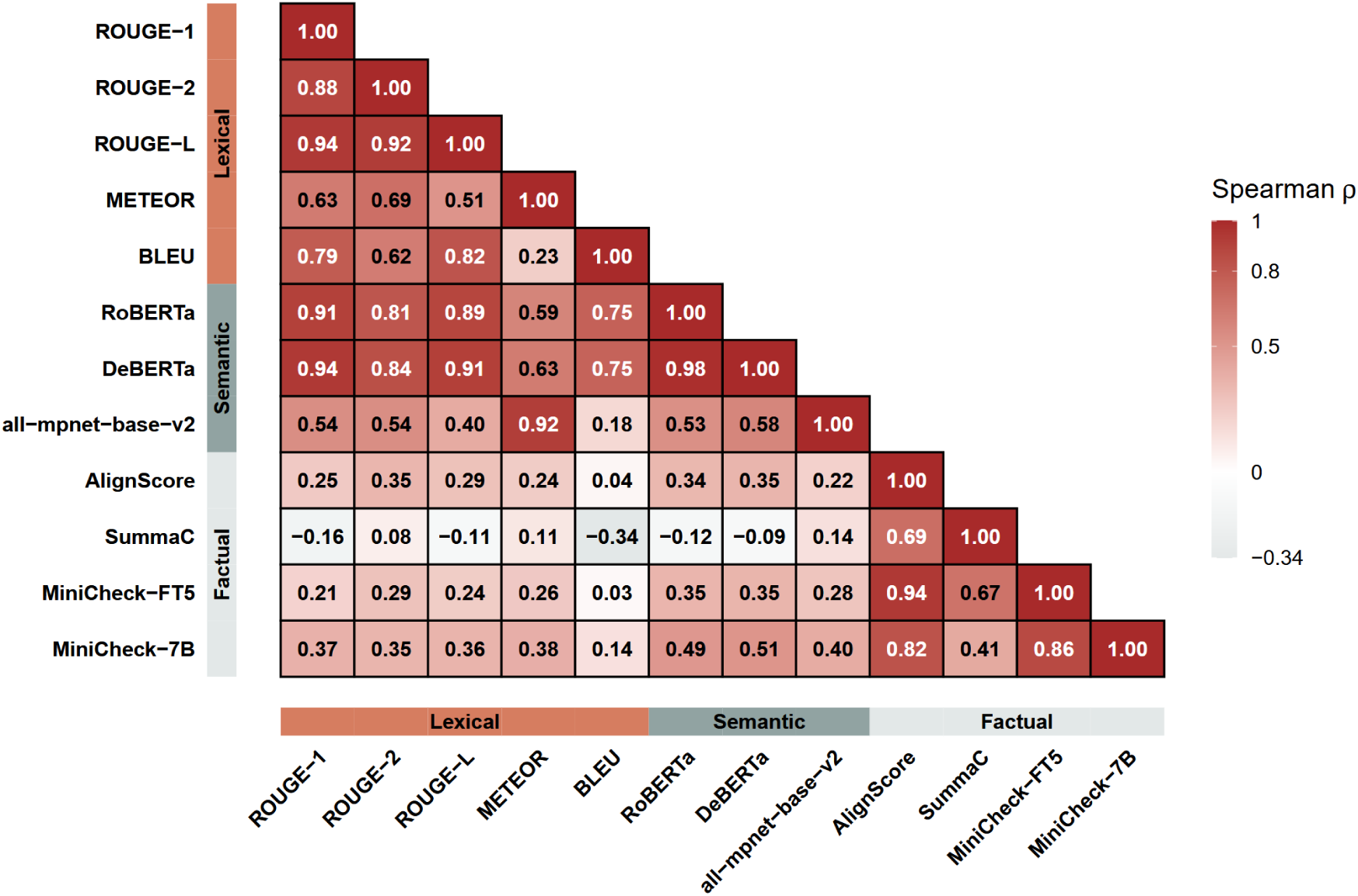
Spearman correlation between evaluation metrics: Spearman correlation coefficients (*ρ*) between two metrics based on their mean scores across all models are given. Metric categories (lexical, semantic, factual) are indicated on the x and y axes. For visualization purposes, cells with values higher than 0.8 are shown with white text.

As the different metric dimensions operate on different value ranges, we applied z-score normalization to the dimension-wise averages across all models, ensuring that lexical, semantic, and factual metrics contribute comparably to the final evaluation.

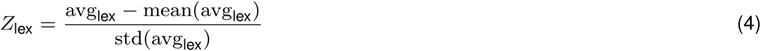

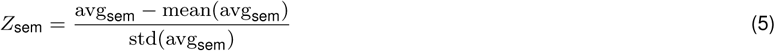

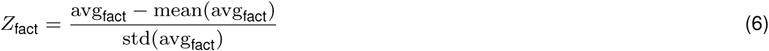

The Overall Performance Score (OPS) was computed as the mean of the normalized dimension scores:

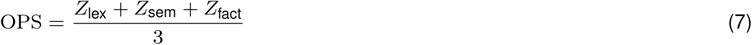

We finally ranked models by overall performance, resulting in the performance rank, with lower ranks indicating better performance. To examine which model performed best in each dimension (lexical, semantic, factual), we additionally computed per-dimension ranks, reported alongside the overall rank.

Differences in overall performance between model categories and families were assessed using Welch’s analysis of variance (ANOVA), followed by pairwise post-hoc comparisons with Games-Howell tests. This procedure is well suited for groups with unequal sample sizes and heterogeneous variances, conditions that apply to our benchmark due to the heterogeneity of models within each category and family. Adjusted *p*-values were used to determine the statistical significance of between-group differences.

### Data Leakage Analysis

To assess whether benchmark articles were part of the models’ pretraining data, we conducted a data leakage analysis by comparing their publication dates against reported model training cutoff dates. If an article was published before a model’s training cutoff date, it may have been included in the model’s internal knowledge, potentially giving the model access to information beyond the input provided (title and abstract).

Publication dates for all benchmark articles were retrieved using the Crossref API. For each model, the reported training cutoff date was collected from official model documentation (i.e., provider-released model descriptions) or, when unavailable, from community-curated sources^50^. In total, 21 cutoff dates were confirmed from primary sources, 9 were obtained from community sources, 17 were estimated based on publicly available information (e.g., Hugging Face documentation or associated publications), and 13 (primarily from newer and less-well documented models) remained unknown. We then compared article publication dates with model cutoff dates to determine the proportion of benchmark articles published on or before each model’s training cutoff. This proportion represents the theoretical maximum share of benchmark articles that could have been present in the model’s training data.

To further investigate models with high potential overlap, we performed an abstract completion probe. Notably, all models included in this abstract completion probe had cutoff dates confirmed from either official or community sources. In this test, models were prompted in completion mode with the first sentence of an article abstract and asked to generate the continuation. For each model, this procedure was done separately for abstracts published before and after the model’s training cutoff date. The similarity between generated summaries and the original abstracts were measured using ROUGE-L scores, as ROUGE-L is based on the longest common subsequence (LCS), which makes it sensitive to long in-order token overlaps. If models had memorized benchmark articles during training, completions for pre-cutoff abstracts would be expected to show significantly higher similarity scores compared to post-cutoff abstracts. Differences in ROUGE-L similarity between pre- and post-cutoff articles were assessed using one-sided Mann-Whitney U tests.

This two-stage analysis provides a practical estimation of whether benchmark articles have influenced model outputs through previous exposure during pretraining.

### Expert Assessment

To assess whether automatic evaluation metrics align with human judgments, we conducted an expert evaluation of the generated summaries. Eight evaluators with professional backgrounds in biomedical research and drug discovery participated. A web-based questionnaire was developed and evaluators were asked to rate generated summaries using the four quality dimensions defined in the SummEval framework^51^, a widely adopted standard for summarization evaluation that has also been used in recent LLM-based evaluation frameworks such as G-Eval^52^. The evaluated dimensions were: coherence (logical structure and organization of the summary), fluency (grammatical correctness and readability), relevance (coverage of important information), and consistency (factual alignment with the reference). Each dimension was rated on a five-point Likert scale ranging from 1 to 5.

The assessment was conducted on summaries generated by the two highest-ranked and the two lowest-ranked models identified in the automated benchmark: mistral-small3.2:24b, mistral-small-2506, bigbird-pegasus-large-pubmed, and mT5 multilingual XLSum. For evaluation, one article was randomly selected from each journal category, resulting in 20 biomedical articles and 80 assessments per expert.

Each evaluator assessed the same set of summaries, but to reduce potential ordering effects, the presentation order was randomized. Model identities were hidden to ensure an unbiased, blinded evaluation. For each paper, the evaluation followed a two-step process. In the first step, evaluators were shown the author-written highlights along with the generated summary and asked to rate coherence, fluency, and relevance. The author-written highlights served as a reference for the key findings, allowing evaluators to assess whether the generated summaries captured the most important information and enabling comparison of human judgments with lexical and semantic evaluation metrics.

In the second step, evaluators were shown the title and abstract of the original article alongside the generated summary to assess factual consistency and detect potential hallucinations. This mirrors the input used for the automatic factuality metrics, enabling direct comparison.

Inter-rater agreement was quantified using Gwet’s AC2 coefficient^53^ with quadratic weights, chosen for its robustness to skewed score distributions and capturing absolute agreement, together with the mean pairwise Spearman’s *ρ*, capturing agreement in relative ranking. Pairwise differences in ratings between models were tested using Mann-Whitney U tests with Bonferroni correction for multiple comparisons.

### LLM-as-a-judge evaluation

We complement the evaluation metrics with an LLM-as-a-judge approach^54^ using three models from distinct providers (Anthropic, OpenAI, Mistral) in a panel-of-evaluators design^55^ to reduce intra-model bias relative to any single judge. We deliberately used mid-tier models while limiting reasoning efforts, rather than using each provider’s flagship model, to prevent overthinking. Each judge emitted a brief rationale before scoring (explain-then-rate, which aids alignment with human ratings^56^), rating each of the four rubric dimensions on a 1 to 5 scale via forced structured outputs and receiving the same evidence as the human experts (title, abstract, reference highlights, and the summary). The panel score for each summary-dimension is the median across the three judges. This test was introduced as a bias-control and robustness measure rather than as a source of additional independent signal (see Limitations). We validated the judge panel against the 80 expert assessments using summary-level Spearman correlation and then applied it to all 62 models on the same 20 publications (one per journal), rather than the full 1,000-document benchmark. All correlations use scipy.stats.spearmanr.

### Benchmarking Framework

The benchmark framework, including all downstream statistical tests and supplementary analyses, was implemented in Python 3.12. Gold standard data were retrieved from open-access articles published by *ScienceDirect* and *Cell Press* through manual extraction of titles, abstracts, and highlights sections, along with metadata including publication URLs, identifiers, section types, and article types where available. Extraction followed an identical procedure for both publishers, and each record was checked for completeness of the required fields. All data were stored in machine-readable JSON format.

The framework was implemented using the Python standard library supplemented by several specialized packages: pandas^57^ for data import and export, scikit-learn^58^ for computing cosine similarities of embeddings and TF-IDF vectors, networkx^59^ for graph construction and the PageRank algorithm^60^. Additional evaluation metrics were computed using NLTK^61^ for METEOR and BLEU scores, ROUGE-score, BERT-score^43^, AlignScore^47^, the summac package implementing the SummaC-ZS model^48^, MiniCheck^49^, and sentence-transformers^62^ with the all-mpnet-base-v2 model^46^. Statistical analyses used pingouin for Welch’s ANOVA and Games-Howell tests, SciPy for Spearman correlations and Mann-Whitney U tests, and a custom implementation of Gwet’s AC2 available in the project repository.

Communication with proprietary closed-source LLMs was facilitated through the official Python APIs provided by Anthropic, Mistral AI, and OpenAI. Local LLM execution was performed on a workstation equipped with a NVIDIA RTX A4000 GPU (16GB VRAM) running Ollama as a backend service, accessed through its Python API along with the transformers library^63^.

All LLMs were configured with a temperature parameter of 0.2 to optimize reproducibility while avoiding completely deterministic outputs. For the latest generation of OpenAI models featuring adaptive reasoning capabilities, the configuration was set to text.verbosity = low and reasoning.effort = minimal. The full set of parameters and prompts are documented in the config.py file in the GitHub repository.

### Data Availability

The source code, documentation, gold standard dataset, and processed results are available at: github.com/Delta4AI/LLMTextSummarizationBenchmark.

## Results

Our benchmark results offer a comparative view of summarization performance across all evaluated models. We first examined the correlations among the chosen evaluation metrics and, based on the findings, identified the best-performing model across lexical, semantic, and factual dimensions. Next, we compared model performance across categories and families to identify significant differences. We additionally performed a data leakage analysis to ensure that inclusion of benchmarking data in training data had no impact on the results. We then analyzed how well automatic metrics aligned with human judgment based on expert level result assessment, and finally assessed whether an LLM-as-a-judge panel could reproduce these rankings.

### Metric Correlations

Pairwise Spearman correlations were computed to analyze the relationships between the different evaluation metrics (Figure 2). Strong positive correlations were observed among most lexical-based metrics (ROUGE-1/2/L, METEOR, and BLEU), with the correlation between METEOR and BLEU marking an exception (*ρ* = 0.23). ROUGE variants showed almost identical behavior (*ρ ≥* 0.88), while BLEU and METEOR demonstrated slightly weaker but still substantial alignment with ROUGE measures (*ρ* = 0.51 *−* 0.82).

Most semantic-based metrics (RoBERTa, DeBERTa, and all-mpnet-base-v2) showed high internal consistency (*ρ >* 0.5), reflecting their shared focus on semantic similarity. When compared with lexical-based metrics, correlations were moderate to strong in most cases, indicating that both categories capture related but not identical dimensions of summary quality.

Concerning factual consistency metrics, AlignScore and the two MiniCheck variants exhibited very strong correlations with each other (*ρ* = 0.82 *−* 0.94), while SummaC showed moderately positive correlations with the other factual metrics (*ρ* = 0.41 *−* 0.69). This likely reflects methodological differences, as SummaC relies on zero-shot natural language inference over sentence pairs, while MiniCheck and AlignScore use models specifically optimized for factual consistency assessment. When compared with lexical- and semantic-based metrics, correlations were generally weaker and, in some cases, even slightly negative. This pattern can largely be attributed to the different point of reference used by factual metrics.

Overall, these relationships demonstrate that the various metrics are broadly consistent while providing complementary perspectives. This supports the use of an aggregated “Overall Performance Score” as a balanced indicator of overall summarization performance.

### Overall Model Performance

Based on overall performance ranks, referred to as “Performance Rank” and derived from our multi-metric evaluation framework (Overall Performance Metric), models from the Mistral family were top-ranked across the three evaluated metric dimensions, as depicted in Figure 3.

**Figure 3:**
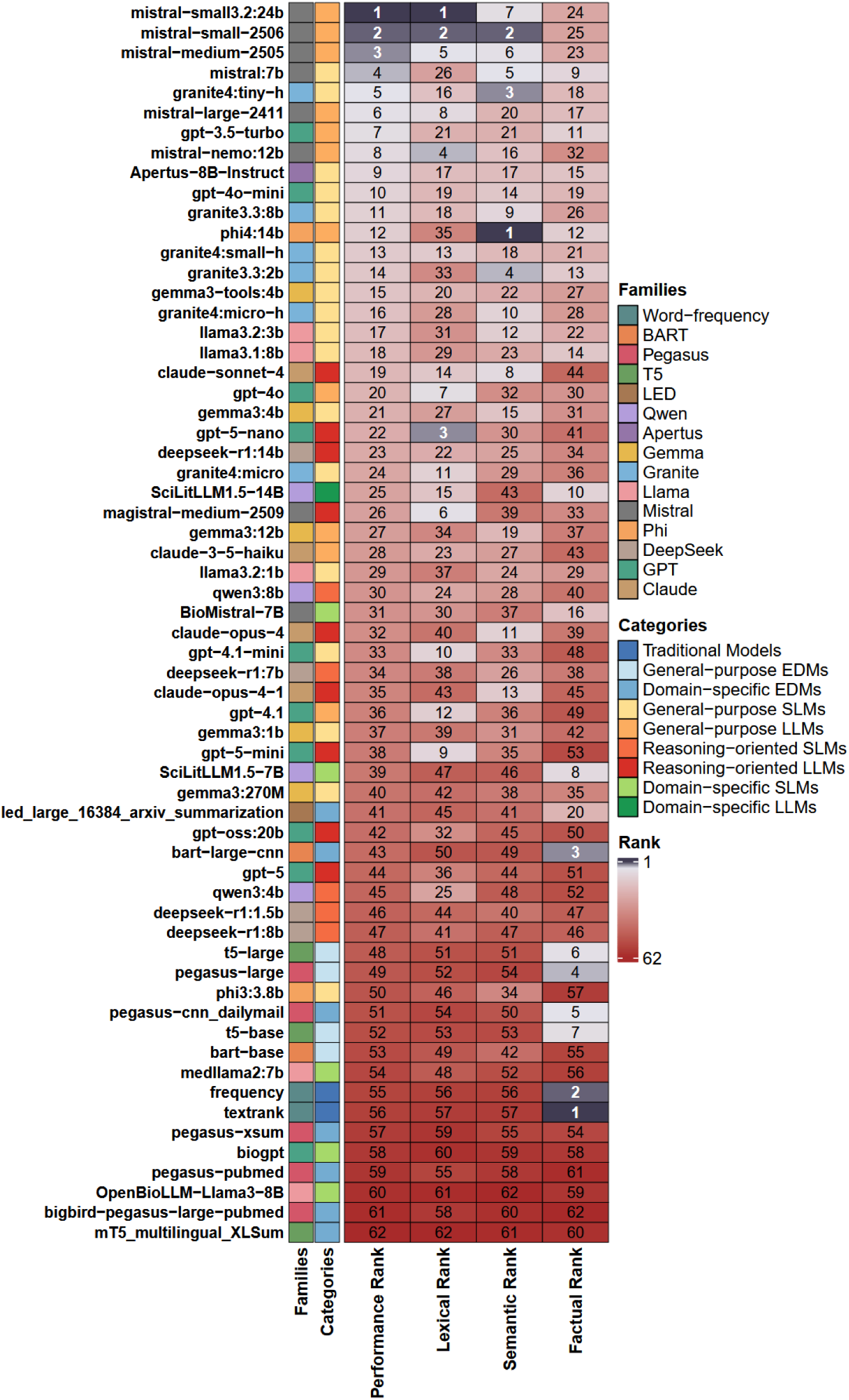
Ranking of summarization models: Overview of the performance of all evaluated models across lexical, semantic, and factual metric dimensions. Each row corresponds to one model. Columns show the overall Performance Rank and independent ranks for each metric dimension, with lower ranks indicating better performance. The figure displays each model’s family and category. Models are sorted by the overall Performance Rank. Ranks are displayed using a continuous diverging color gradient, ranging from navy-blue tones for top-performing models through light neutral shades for mid-ranked models to dark red tones for low-performing models.

The mistral-small3.2:24b model ranked first, followed by mistral-small-2506 and mistral-medium-2505. The lowest-ranked models included OpenBioLLM-Llama3-8B, bigbird-pegasus-large-pubmed and pegasus-pubmed from the PEGASUS family, and mT5 multilingual XLSum from the T5 family. Overall, domain-specific SLMs and EDM models showed poor performance across all the different dimensions of metrics.

Among the ten top-ranked models, six were general-purpose LLMs and four were general-purpose SLMs. A similar trend was observed in the top half of the ranking (positions 1 to 31) with the presence of just a few reasoning-oriented and domain-specific models. In contrast, in the lower half of the ranking, where models performed poorly across most metric dimensions, the majority were reasoning-oriented SLMs, general-purpose EDMs, domain-specific EDMs, and traditional models.

Considering each metric dimension separately, several Mistral models ranked high in the lexical dimension, with GPT-5-nano, a reasoning-oriented LLM from the GPT family, ranking third. In the semantic dimension, Mistral models achieved high ranks, but the highest positions were occupied by the Phi4:14b general-purpose LLM and several Granite models, including granite4:tiny-h and granite3.3:2b. Finally, the highest ranks based on the factual dimension were covered by traditional models and some EDMs. Detailed performance indications for each model across all individual ranked metrics are provided in supplementary Figure 8.

Models could also decline to summarize by returning the INSUFFICIENT FINDINGS token. This was rare overall, occurring in 736 of 62,000 generated summaries (1.2%) and exclusively among prompt-based models. Usage was concentrated in the two domain-specific biomedical models OpenBioLLM-Llama3-8B (20.0%) and medllama2 (9.7%), while all other models returned it in fewer than 6% of cases.

### Category Comparisons

To compare model performance across categories, we displayed the distribution of overall performance scores within each category, along with the best- and worst-performing individual models (Figure 4a). Results of the Games-Howell post-hoc comparisons between individual groups are given in Figure 4b.

**Figure 4:**
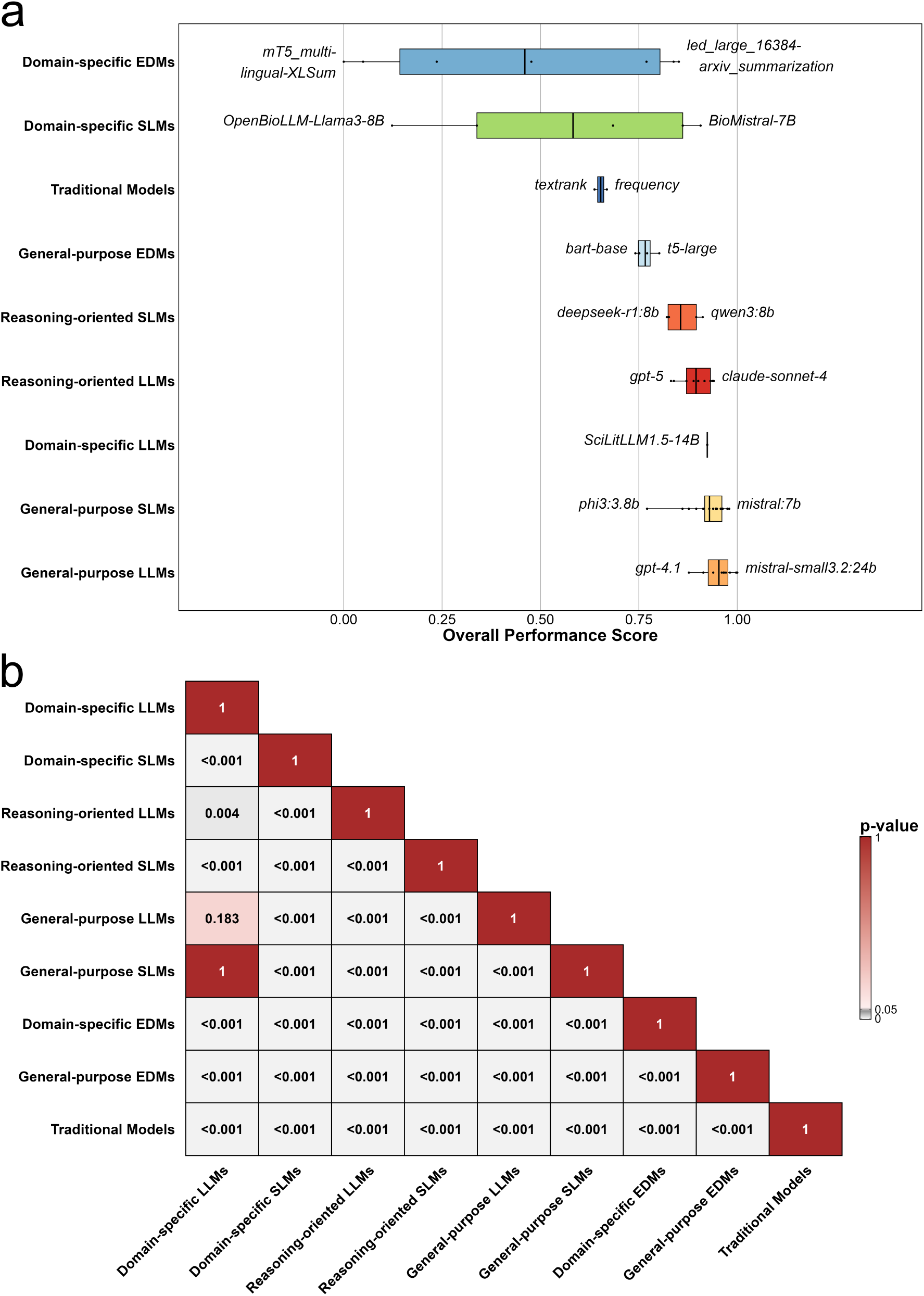
(a) Overall performance of model categories: Distributions of overall performance scores across the nine model categories, with each model represented by a black dot. Categories are ordered from lowest to highest-performing according to the median performance score. The best- and worst-performing models of each category are highlighted. **(b) Games-Howell Post-hoc Test across model categories:** Games–Howell pairwise adjusted *p*-value matrix comparing all category pairs. Each cell shows the adjusted *p*-value for the difference in metric mean scores between two categories, with white-to-gray shading indicating smaller *p*-values.

All general-purpose LLMs achieved high overall scores, thus forming the top category of models, with general-purpose SLMs following second, however showing a slightly wider spread in performance scores. Domain-specific LLMs and reasoning-oriented LLMs followed in third and fourth place based on average overall performance.

Domain-specific EDMs, domain-specific SLMs, traditional models, and general-purpose EDMs performed significantly worse as compared to general-purpose language models (LMs). Domain-specific EDMs and domain-specific SLMs displayed the widest spread of performance (Figure 4a).

### Family Comparisons

To complement the category-level findings, we examined performance across model families, since models within the same family often share architectural features or training strategies that may cause consistent performance trends. Figure 5a shows the distribution of overall performance scores across all model families. Similar to the category-level analysis, there were clear performance differences between architectural lineages. The Mistral family achieved the strongest overall results, with several members ranking among the top performers in the benchmark. Families dominated by modern LLM or SLM architectures, such as Granite, Claude, and Gemma, consistently outperformed more traditional extractive approaches and families built on encoder-decoder architectures such as longformer encoder-decoder (LED), BART, T5 and PEGASUS.

**Figure 5:**
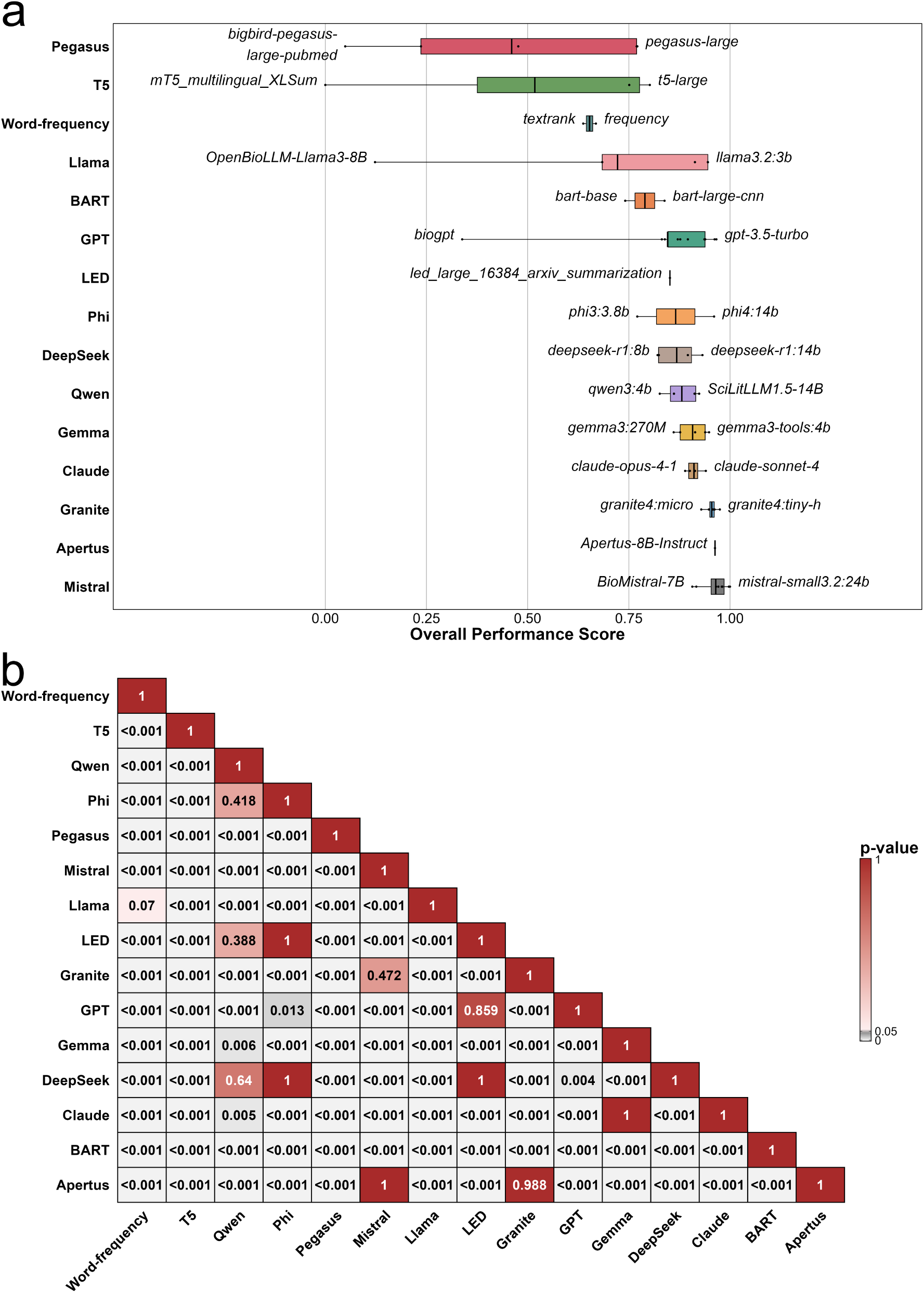
(a) Overall performance of model families: Distributions of overall performance scores across all model families, with each model represented by a black dot. Families are ordered from lowest to highest-performing according to the performance score. The best- and worst-performing models of each family are highlighted. **(b) Games-Howell Post-hoc Test across model families:** Games–Howell pairwise adjusted *p*-value matrix comparing all family pairs. Each cell shows the adjusted *p*-value for the difference in metric mean scores between two families, with white-to-gray shading indicating smaller *p*-values.

In particular, the GPT and Llama families showed lower performance scores than expected given the competitive performance of their top models. This was primarily due to the inclusion of domain-specific small models, such as BioGPT and OpenBioLLM-Llama3-8B, which performed substantially worse than their general-purpose counterparts and therefore pulled down the family-level averages.

Regarding the Games-Howell post-hoc matrix in the family comparison (Figure 5b), a combination of significant and non-significant differences between families was observed, reflecting the greater heterogeneity within some lineages. While many high-performing families differed significantly from traditional and encoder-decoder dominated families, several comparisons among modern SLMs and LLMs dominated families did not reach statistical significance.

### Data Leakage Assessment

#### Cutoff Overlap Results

Across the evaluated models, potential overlap ranged from minimal levels of 5% for earlier models with training cutoffs prior to 2024, up to 40.2% for more recent LLMs with reported cutoff dates in early 2025. Most models showed low overlap (*≤* 20%), indicating that the majority of benchmark articles were published after their respective training cutoff dates. A subset of eight models, including recent versions of Claude, Deepseek, and one GPT model, exhibited moderate levels of potential overlap (24 *−* 40.2%) and were further analyzed via an abstract completion probe in the next step.

A complete overview of training cutoff dates and corresponding potential overlap values for all models is provided in Supplementary Table 8.

#### Abstract Completion Probe

To assess whether high potential overlap translated into memorization effects, we conducted an abstract completion probe for the eight models showing moderate levels of potential overlap. For each model ROUGE-L scores between generated completions and the original abstracts were compared for articles published before and after the respective training cutoff dates.

Average ROUGE-L scores were nearly identical for pre-cutoff and post-cutoff articles (both at approximately 13.5%) and there were only negligible differences at the individual model level (Table 3). Statistical testing using a one-sided Mann-Whitney U test revealed no significant increase in similarity for pre-cutoff articles in any of the tested models. Effect sizes were consistently close to zero, which further indicates that there is no systematic difference between the two groups.

**Table 3:** Abstract completion probe results. Shows ROUGE-L scores for completions of abstracts published before (pre-cutoff) and after (post-cutoff) the respective training cutoff dates. Δ denotes the difference between both groups. Statistical significance was assessed using a one-sided Mann-Whitney U test.

| Model | Cutoff | Pre ROUGE-L | Post ROUGE-L | $\Delta$ | 95% CI | *p*-value | Effect(*r*) |
| --- | --- | --- | --- | --- | --- | --- | --- |
| claude-sonnet-4-20250514 (closed-source) | 2025-03 | 15.2 | 15.2 | +0.0 | [-0.5, +0.4] | 0.598 | +0.01 |
| claude-opus-4-20250514 (closed-source) | 2025-03 | 15.3 | 15.0 | +0.2 | [-0.3, +0.8] | 0.239 | -0.03 |
| claude-opus-4-1-20250805 (closed-source) | 2025-03 | 15.2 | 14.9 | +0.3 | [-0.2, +0.8] | 0.161 | -0.04 |
| deepseek-r1:1.5b | 2025-01 | 10.3 | 10.5 | -0.1 | [-0.7, +0.4] | 0.598 | +0.01 |
| deepseek-r1:7b | 2025-01 | 11.7 | 11.9 | -0.2 | [-0.6, +0.3] | 0.524 | +0.00 |
| deepseek-r1:8b | 2025-01 | 13.2 | 13.5 | -0.3 | [-0.7, +0.2] | 0.834 | +0.04 |
| deepseek-r1:14b | 2025-01 | 13.4 | 13.4 | -0.1 | [-0.4, +0.3] | 0.662 | +0.02 |
| gpt-5-2025-08-07 (closed-source) | 2024-10 | 13.7 | 13.5 | +0.2 | [-0.2, +0.7] | 0.399 | -0.01 |

If memorization effects had occurred, substantially higher similarity scores would be expected for abstracts from pre-cutoff articles. As this was not the case, we concluded that the models were not reproducing memorized content from their training data.

### Expert Assessment Results

The results of the expert assessment are summarized in Figure 6 and Table 4. Values are reported as mean *±* standard error of the mean (SEM). The two Mistral-based models (mistral-small3.2:24b and mistral-small-2506) consistently outperformed the two EDMs (bigbird-pegasus-large-pubmed and mT5 multilingual XLSum) across all four evaluation dimensions.

**Figure 6:**
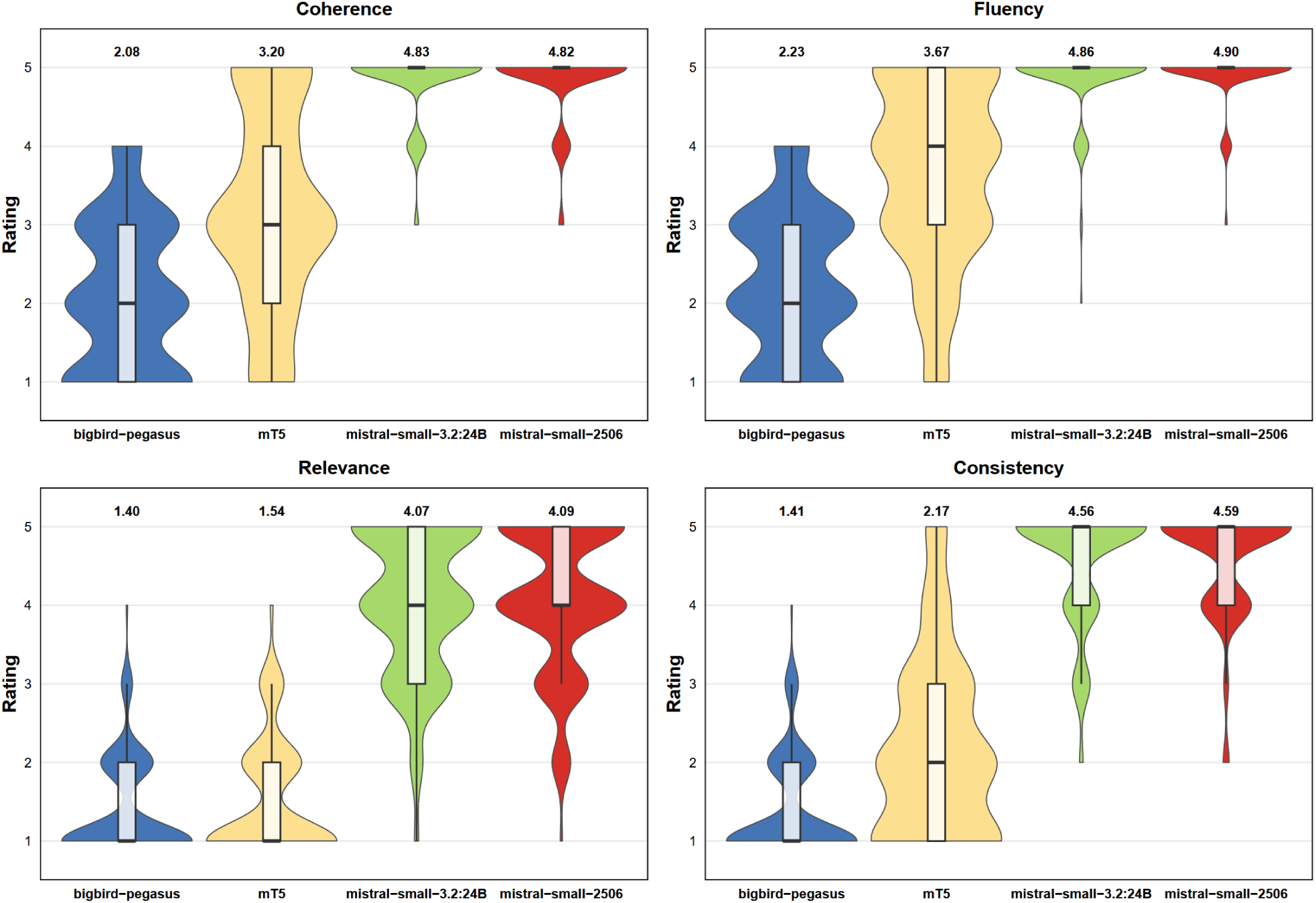
Expert assessment results across four quality dimensions: Distribution of expert ratings for coherence, fluency, relevance, and consistency for each model are shown with average ratings shown above the box-violin plots.

**Table 4:** Expert assessment results (mean. *±* **SEM,** *n***=160 per model):** Gwet’s AC2 (quadratic weights) and mean pairwise Spearman’s *ρ* measure inter-rater agreement.

| Model | Coherence | Fluency | Relevance | Consistency |
| --- | --- | --- | --- | --- |
| BigBird-Pegasus | 2.08 ± 0.08 | 2.23 ± 0.07 | 1.40 ± 0.05 | 1.41 ± 0.05 |
| mT5-XLSum | 3.20 ± 0.10 | 3.67 ± 0.09 | 1.54 ± 0.06 | 2.17 ± 0.09 |
| Mistral-Small-3.2 (24B) | 4.83 ± 0.03 | 4.86 ± 0.03 | 4.07 ± 0.07 | 4.56 ± 0.06 |
| Mistral-Small-2506 | 4.82 ± 0.04 | 4.90 ± 0.03 | 4.09 ± 0.07 | 4.59 ± 0.05 |
| Gwet’s AC2 | 0.735 | 0.794 | 0.713 | 0.735 |
| Spearman’s $\rho$ (mean) | 0.707 | 0.722 | 0.799 | 0.801 |

Overall AC2 = 0.725, Overall $\rho$ = 0.765

For coherence and fluency, both Mistral models scored substantially higher than the EDMs. In coherence, they reached scores of 4.83 *±* 0.03 and 4.82 *±* 0.04, compared with 3.20 *±* 0.10 for mT5 and 2.08 *±* 0.08 for bigbird-pegasus. The same pattern was observed for fluency, where the Mistral models achieved scores of 4.86 *±* 0.03 and 4.90 *±* 0.03, clearly exceeding mT5 (3.67 *±* 0.09) and bigbird-pegasus (2.23 *±* 0.07). All pairwise comparisons between Mistral models and EDMs were statistically significant (all *p <* 0.001 after Bonferroni correction). The two EDMs also differed significantly from each other in both dimensions, while no significant differences were found between the two Mistral models.

Differences were even more pronounced for relevance and consistency. The Mistral models again achieved high scores in relevance (4.07 *±* 0.07 and 4.09 *±* 0.07), while mT5 (1.54 *±* 0.06) and bigbird-pegasus (1.40 *±* 0.05) scored markedly lower. A similar pattern was seen for consistency, where the Mistral models reached scores of 4.56 *±* 0.06 and 4.59 *±* 0.05, compared to 2.17 *±* 0.09 for mT5 and 1.41 *±* 0.05 for bigbird-pegasus. Again, all Mistral versus EDM comparisons were statistically significant (*p <* 0.001). The EDMs differed significantly in consistency, but not in relevance (*p* = 0.7308), and no significant differences were found between the two Mistral models.

Inter-rater agreement was strong on both measures. Gwet’s AC2 indicated substantial absolute agreement across all dimensions, ranging from 0.713 (relevance) to 0.794 (fluency), with an overall value of AC2 = 0.725. Mean pairwise Spearman correlations were likewise strong across all dimensions, ranging from *ρ* = 0.707 (coherence) to *ρ* = 0.801 (consistency), with an overall mean of *ρ* = 0.765 (all *p <* 0.001). These results show that evaluators consistently agreed both on the absolute quality of the summaries and on the relative ranking between models.

### LLM-as-a-judge evaluation

The LLM-as-a-judge panel agreed closely with expert assessments on every dimension (Table 5). To benchmark against human reliability, we computed a leave-one-out ceiling on the same target the judges were scored against: each expert’s ratings were correlated against the mean of the remaining experts. The judge’s mean agreement (*ρ* = 0.87) exceeded this ceiling (*ρ* = 0.82), leading to the assumption that it tracked the expert consensus at least as closely as a typical individual expert.

**Table 5:**
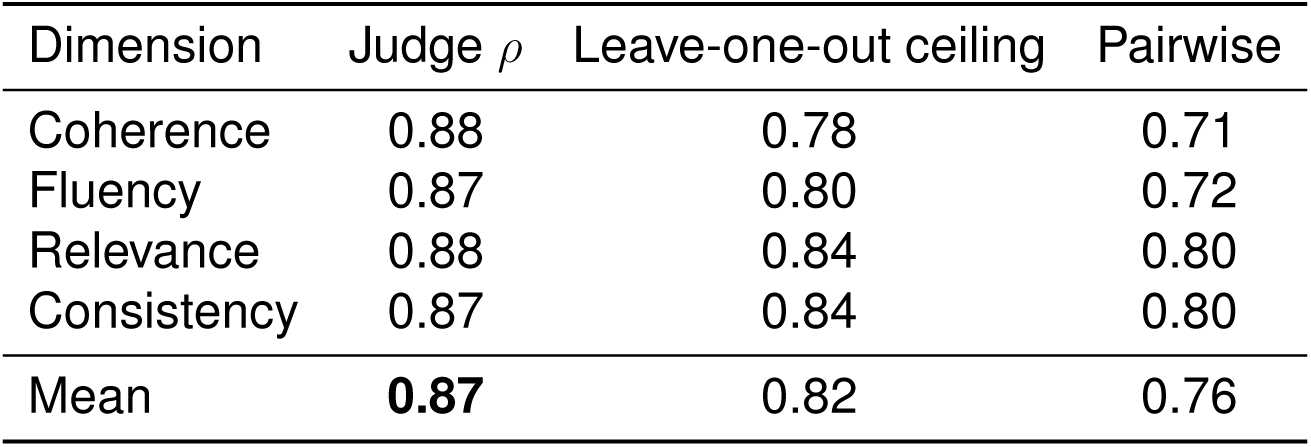
LLM-as-a-judge panel vs. expert assessments (Spearman’s *ρ*, *n* = 80 summaries). *Diagonal* : judge dimension vs. the same expert dimension. *Leave-one-out ceiling*: each expert vs. the mean of the others (fair, same-target human upper bound). *Pairwise*: raw inter-rater agreement between two individual experts.

| Dimension | Judge $\rho$ | Leave-one-out ceiling | Pairwise |
| --- | --- | --- | --- |
| Coherence | 0.88 | 0.78 | 0.71 |
| Fluency | 0.87 | 0.80 | 0.72 |
| Relevance | 0.88 | 0.84 | 0.80 |
| Consistency | 0.87 | 0.84 | 0.80 |
| Mean | **0.87** | 0.82 | 0.76 |

The judge panel did not separate the four dimensions well. Its agreement with the matching expert dimension (diagonal *ρ* = 0.87) was barely higher than its agreement with the other dimensions (off-diagonal *ρ* = 0.85), a gap of only 0.02. In effect the judge measured a single overall-quality signal rather than four independent constructs, so its per-dimension scores should not be over-interpreted.

We questioned whether the LLM-as-a-judge panel was capable of ranking the 62 models in a comparable manner to the evaluation metrics (Table 6). The judge ranking agreed strongly with the lexical, semantic, and composite performance ranks (*ρ* = 0.75-0.83). The factual metrics were an exception (*ρ* = 0.09, n.s.), as they reward verbatim copying, so extractive systems (TextRank, frequency, T5, PEGASUS) ranked highly on factuality but performed poorly under the judge - a construct difference rather than a judge failure.

**Table 6:** System-level rank concordance between the LLM-as-a-judge ranks and each evaluation metric category across all *n* = 62 models. Judge ranks derive from the 20-publication subset, metric ranks use the full benchmark. This measures agreement between two automatic methods, not human validation.

| Metric family | $\rho$ | $\tau$ | $p$ |
| --- | --- | --- | --- |
| Lexical | 0.75 | 0.57 | $< 10^{-11}$ |
| Semantic | 0.83 | 0.64 | $< 10^{-15}$ |
| Performance | 0.78 | 0.61 | $< 10^{-12}$ |
| Factual | 0.09 | 0.06 | 0.48 (n.s.) |

To analyze the tendency of judges to favor their own model family, we computed, for each judge, how far its score sat above or below the mean of the other two judges, and contrasted its own-family models with all others (Table 7). Anthropic exhibited a clear self-preference effect (+0.30 on the 1-5 scale), whereas OpenAI and Mistral showed no detectable differential effect (*≤* 0.08 in magnitude). The direction is consistent with self-preference reported for rubric-based judging^64,65^. Taking the median across the three judges damped this bias in the final scores.

**Table 7:** LLM-as-a-judge self-preference: a judge’s mean score for its own model family minus its score for other systems, each measured relative to the other two judges (1-5 scale). Positive values indicate inflation of the judge’s own family.

| Judge provider | Self-preference effect |
| --- | --- |
| Anthropic | +0.30 |
| OpenAI | −0.08 |
| Mistral | −0.02 |

## Discussion

Our benchmarking analysis of 62 text summarization models revealed clear performance differences. General-purpose LLMs achieved the highest summarization quality across all metric dimensions, closely followed by general-purpose SLMs, domain-specific LLMs, and reasoning-oriented LLMs. In contrast, domain-specific SLMs, encoder–decoder architectures, and traditional extractive methods performed significantly worse regarding overall performance. These results highlight the clear progression from extractive and encoder–decoder approaches toward transformer-based models, while also showing that domain-specific fine-tuning alone does not necessarily lead to improved summarization quality.

Comparing models by architecture, size, and domain focus shows that, overall, LLMs perform best, likely due to their large number of parameters, which enable them to better understand the complex context typical for biomedical literature. While lightweight models like Gemma3:270M may lack the capacity to handle this complexity, SLMs remain competitive, with some models, such as mistral:7b and granite4:tiny-h, even outperforming several LLMs across the three metric dimensions. This may be attributed to the fact that smaller datasets are often more curated and of higher quality compared to the large amount of data required to train a large model^66^.

Interestingly, medium-sized models (e.g., many models in the Mistral family) outperform larger proprietary ones. These models seem to reach an optimal balance between parameter count and performance, where additional parameters could disrupt this equilibrium, potentially leading to over-fitting or plateaus^67^.

We also found that overall general-purpose models outperform both domain-specific models specialized in the biomedical domain and those specialized for text summarization, regardless of size. A possible explanation might be that domain-specific models fine-tuned on biomedical text might be better for learning and understanding the complex biomedical terminology or lexical patterns but fail in summarization tasks. On the other hand, models specifically designed for text summarization might be good at summarizing in general but fail at capturing the complex biomedical meaning. Therefore, generalist models, leveraging their broad knowledge, seem to perform better^68^. Additionally, domain-specific models can “forget” the general knowledge acquired during the pre-training phase, experiencing a phenomenon called “catastrophic forgetting”, which represents an issue when the task requires both biomedical knowledge and context understanding for text summarization^69^.

Most reasoning-oriented models ranked in the middle, indicating moderate performance. This likely reflects their multi-step reasoning nature, which can be advantageous for tasks that require breaking down problems into sequential steps, such as mathematical operations or computer programming, but seems less suited for text summarization, which requires semantic compression and factual grounding^70^.

Traditional models ranked highest on the factual dimension, likely because their extractive approach generates summaries closely aligned with the source abstracts, which factuality metrics reward by design^71^.

The expert assessment supported the findings of the automated benchmark. The two highest-ranked models consistently outperformed the two lowest-ranked models across all four evaluation dimensions. This alignment suggests that the automatic metrics capture relevant aspects of summary quality and are broadly consistent with human judgments.

The results of this benchmark provide useful guidance for selecting summarization models in biomedical and scientific settings. The strong performance of general-purpose LMs indicates that broad, diverse pretraining is often more advantageous than narrow domain adaptation when dealing with unseen scientific content.

Another key consideration is the trade-off between output quality and processing efficiency. While LLMs achieved the highest overall scores, SLMs deliver competitive results at substantially lower computational cost^72^, which makes them especially attractive for large-scale or resource-constrained applications. Choosing between large and small models therefore depends not only on desired output quality but also on the intended scale of summarization.

Overall, the findings suggest that general-purpose LMs currently offer the most reliable and practical choice for biomedical text summarization. Their consistent performance across evaluation criteria demonstrates that broad generalization outweighs the marginal gains from more narrowly specialized or fine-tuned approaches, many of which are not primarily optimized for text summarization.

### Limitations of the study

Even though our results are based on a robust evaluation framework, several limitations of our study warrant discussion. Model access methods varied across the evaluation due to differing application programming interface (API) capabilities and requirements. Hugging Face models were accessed through their supported interfaces: the pipeline API with task=”summarization” where available, or chat/completion formats for models that did not support the pipeline approach. Ollama models required use of the generate endpoint with merged prompts, while OpenAI, Anthropic, and Mistral models each mandated their respective provider-specific APIs (responses.create, messages.create, and chat.complete) with distinct message structures. We applied hyperparameter normalization where possible, though API-level constraints prevented full standardization. For example, GPT-5 does not support temperature control, instead offering only reasoning-specific parameters. Additionally, proprietary middleware layers may transform requests and responses in undocumented ways, potentially affecting outputs independently of the underlying model architectures. These necessary methodological variations warrant consideration when interpreting performance differences across models. Furthermore, all models received an identical summarization prompt. While this ensures a controlled comparison, it may disadvantage models whose instruction tuning differs from the chosen phrasing. Per-model prompt optimization (e.g. via DSPy^73,74^) could yield different absolute scores and rankings. Our results therefore reflect performance under a fixed instruction, not each model’s attainable ceiling.

Reference-based evaluation cannot fully capture every valid way a good summary might be written, as summarization is largely a subjective exercise. To account for this, we tried to capture summary quality along diverse, complementary dimensions by combining lexical, semantic, and factual metrics, and as a further assessment included a blinded expert evaluation as an independent check on the automatic scores. Nonetheless, since automatic metrics and human ratings have not fully aligned in other settings^5,6^, we cannot formally guarantee that our metrics capture the full space of valid summaries, which remains an important direction for future work.

Regarding our LLM-as-a-judge results, it is notable that the three judges are not independent: correlated errors mean the panel yields far fewer effective votes than its size suggests, and a strong single judge can match it^75^. We therefore treat the panel median as bias-control, not a route to higher accuracy, and its edge over the human leave-one-out ceiling reflects median smoothing rather than above-human reliability. Furthermore, the summary-level agreement pools models of very different quality. Within-model agreement is much lower, so the judge ranks models reliably but separates similar-quality summaries weakly. Finally, the system-level agreement compares two automatic methods, not validation against humans (infeasible with only four expert-rated models).

Another limitation of this benchmarking study is that we focused on a single summarization task: generating concise summaries from biomedical abstracts. This setup provides a clear and well-defined evaluation framework, but the findings may not fully extend to other forms of scientific or biomedical summarization, including full-text articles, clinical trial data, or lay-oriented summaries. Given the rapid evolution of LLMs, these results just capture a specific snapshot in time and may change as newer architectures and models become available.

A related consideration is our choice of reference summary. We used author-written highlights as the reference against which summaries were evaluated, but condensing an abstract (let alone a full publication) into a few bullet points is inherently subjective, and highlights inevitably emphasize some findings over others. We nonetheless regard them as the most suitable reference available, since they are author-generated, standardized across both publishers, and well matched to our target of concise summaries within limited length.

Finally, although we deliberately drew articles from 20 journals spanning a broad range of biomedical and translational fields, our corpus inevitably covers only part of this very diverse domain. We cannot rule out that summarization performance, and the resulting model rankings, would differ for subfields not well presented in our selection, leaving open how well these results generalize across the biomedical domain.

## Conclusion

This benchmark provides a comprehensive evaluation of 62 text summarization methods on a dataset of 1,000 biomedical abstracts paired with standardized highlights sections. Across all lexical overlap, semantic similarity, and factual consistency metrics, general-purpose LLMs were the best-performing models. Their ability to incorporate broad pretraining with strong semantic understanding enabled them to outperform reasoning-oriented models, domain-specific models, and encoder-decoder architectures. Especially medium-sized models achieved the highest overall rankings, which indicates that increased scale beyond this range does not automatically translate to improved summarization quality for biomedical text.

SLMs remain competitive for settings where computational resources are limited as they offer balance between efficiency and output quality. In contrast, domain-specific models did not show a systematic advantage, showing that narrow fine-tuning alone is insufficient to match the generalization of broadly trained LLMs for this task.

Overall, this benchmark provides a systematic reference point for selecting summarization models in biomedical research. By assessing a broad spectrum of architectures under a unified framework we underline the strengths and limitations of currently available systems and highlight the superiority of general-purpose LLMs for scientific text summarization. These insights can support researchers and practitioners in choosing models that balance output quality, computational cost, and practical usability in biomedical workflows.

## Supplemental information

**Figure 7:**
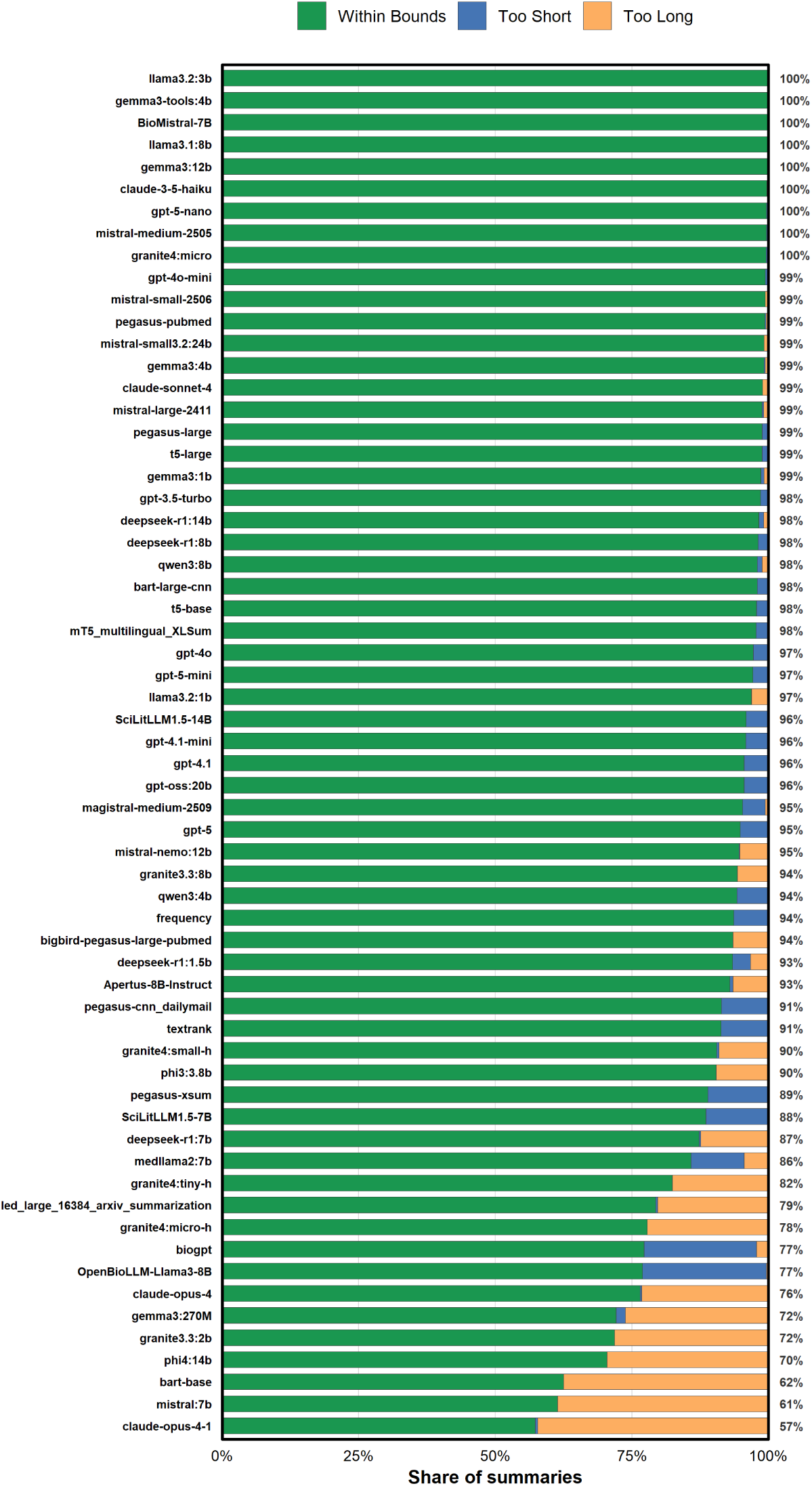
Summary length compliance per method: Proportion of generated summaries that fell within the target length of 15–100 words, were too short, or were too long, shown for each of the 62 evaluated methods. Methods are ordered by decreasing share of within-bounds summaries, with the within-bounds percentage annotated at the right of each bar.

**Figure 8:**
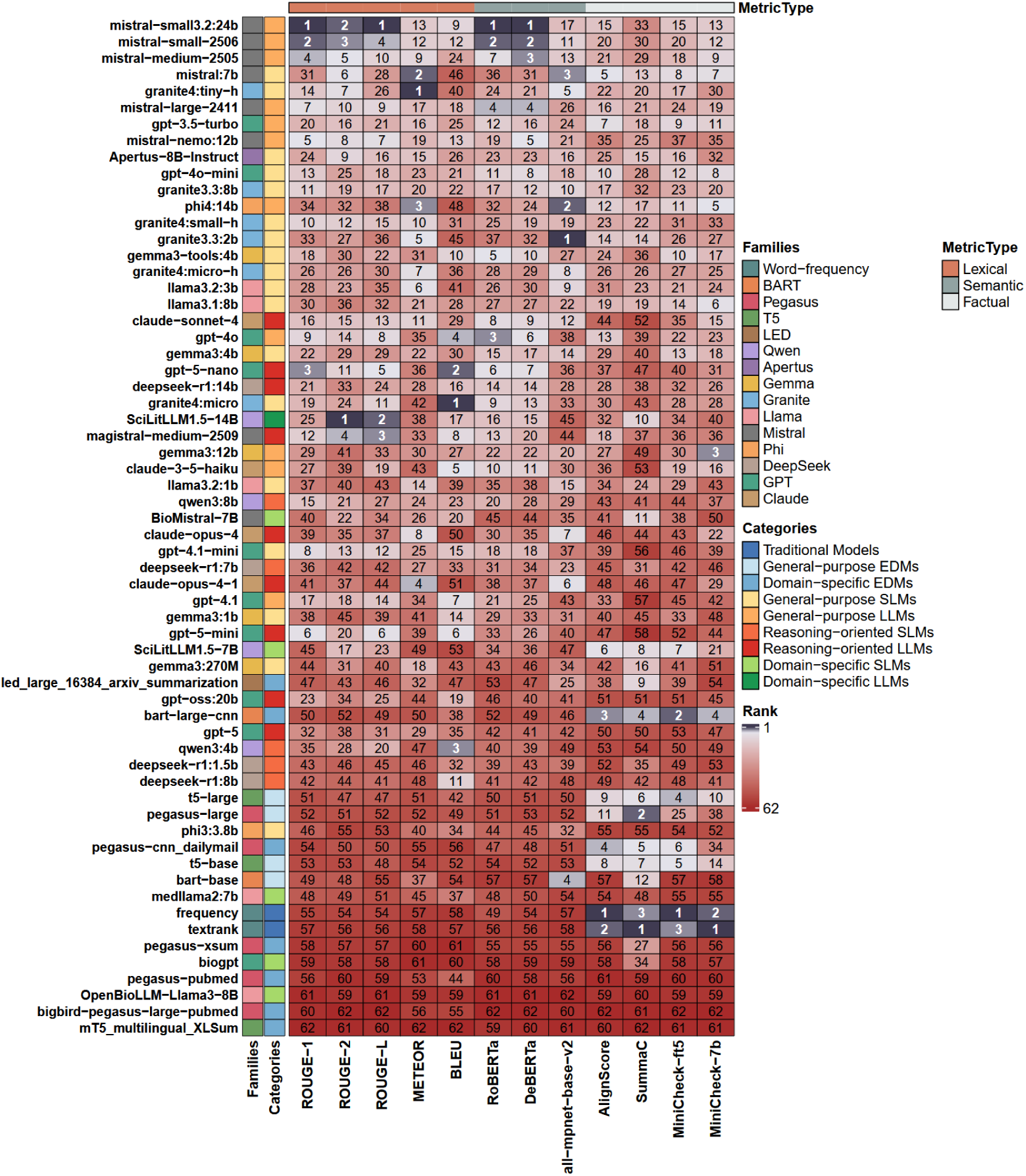
Model performance across metrics: Overview of the performance of all evaluated models across all metrics.

**Table 8:** Training cutoff dates and potential overlap for all evaluated models: The training cutoff indicates the latest point in time up to which a model may have been trained; the source of this information is given in a separate column (confirmed, community, estimated, or unknown). Potential overlap describes the percentage of benchmark articles published on or before the respective cutoff date and therefore represents the theoretical maximum share of articles that could have been present in the model’s training data. Closed-source models are marked accordingly. Models are ordered by decreasing potential overlap.

| Model | Training cutoff | Source | Potential overlap (%) |
| --- | --- | --- | --- |
| claude-sonnet-4-20250514 (closed-source) | 2025-03 | confirmed | 40.2 |
| claude-opus-4-20250514 (closed-source) | 2025-03 | confirmed | 40.2 |
| claude-opus-4-1-20250805 (closed-source) | 2025-03 | community | 40.2 |
| deepseek-r1:1.5b | 2025-01 | community | 33.1 |
| deepseek-r1:7b | 2025-01 | community | 33.1 |
| deepseek-r1:8b | 2025-01 | community | 33.1 |
| deepseek-r1:14b | 2025-01 | community | 33.1 |
| gpt-5-2025-08-07 (closed-source) | 2024-10 | community | 24.0 |
| gemma3:270M | 2024-08 | confirmed | 20.3 |
| gemma3:1b | 2024-08 | confirmed | 20.3 |
| gemma3:4b | 2024-08 | confirmed | 20.3 |
| gemma3:12b | 2024-08 | confirmed | 20.3 |
| PetrosStav/gemma3-tools:4b | 2024-08 | confirmed | 20.3 |
| phi4:14b | 2024-06 | confirmed | 13.8 |
| gpt-oss:20b | 2024-06 | community | 13.8 |
| gpt-4.1 (closed-source) | 2024-06 | confirmed | 13.8 |
| gpt-4.1-mini (closed-source) | 2024-06 | confirmed | 13.8 |
| gpt-5-nano-2025-08-07 (closed-source) | 2024-05 | community | 11.8 |
| gpt-5-mini-2025-08-07 (closed-source) | 2024-05 | community | 11.8 |
| granite3.3:2b | 2024-04 | estimated | 10.3 |
| granite3.3:8b | 2024-04 | estimated | 10.3 |
| claude-3-5-haiku-20241022 (closed-source) | 2024-04 | confirmed | 10.3 |
| swiss-ai/Apertus-8B-Instruct-2509 | 2024-03 | confirmed | 8.5 |
| Uni-SMART/SciLitLLM1.5-7B | 2024 | estimated | 6.6 |
| Uni-SMART/SciLitLLM1.5-14B | 2024 | estimated | 6.6 |
| facebook/bart-large-cnn | 2019 | estimated | 5.0 |
| facebook/bart-base | 2019 | estimated | 5.0 |
| google-t5/t5-base | 2019-04 | estimated | 5.0 |
| google-t5/t5-large | 2019-04 | estimated | 5.0 |
| csebuetnlp/mT5_multilingual_XLSum | 2020 | estimated | 5.0 |
| google/pegasus-xsum | 2020 | estimated | 5.0 |
| google/pegasus-large | 2020 | estimated | 5.0 |
| google/pegasus-cnn_dailymail | 2020 | estimated | 5.0 |
| AlgorithmicResearchGroup/led_large_16384_arxiv_summarization | 2020 | estimated | 5.0 |
| google/pegasus-pubmed | 2020 | estimated | 5.0 |
| google/bigbird-pegasus-large-pubmed | 2020 | estimated | 5.0 |
| microsoft/biogpt | 2021 | estimated | 5.0 |
| aaditya/OpenBioLLM-Llama3-8B | 2023-03 | confirmed | 5.0 |
| BioMistral/BioMistral-7B | 2023 | estimated | 5.0 |
| llama3.1:8b | 2023-12 | confirmed | 5.0 |
| llama3.2:1b | 2023-12 | confirmed | 5.0 |
| llama3.2:3b | 2023-12 | confirmed | 5.0 |
| medllama2:7b | 2022-09 | confirmed | 5.0 |
| phi3:3.8b | 2023-10 | confirmed | 5.0 |
| gpt-3.5-turbo (closed-source) | 2021-09 | confirmed | 5.0 |
| gpt-4o (closed-source) | 2023-10 | confirmed | 5.0 |
| gpt-4o-mini (closed-source) | 2023-10 | confirmed | 5.0 |
| mistral:7b | — | unknown | — |
| mistral-nemo:12b | — | unknown | — |
| mistral-small3.2:24b | — | unknown | — |
| qwen3:4b | — | unknown | — |
| qwen3:8b | — | unknown | — |
| granite4:micro | — | unknown | — |
| granite4:micro-h | — | unknown | — |
| granite4:tiny-h | — | unknown | — |
| granite4:small-h | — | unknown | — |
| mistral-medium-2505 (closed-source) | — | unknown | — |
| magistral-medium-2509 (closed-source) | — | unknown | — |
| mistral-large-2411 (closed-source) | — | unknown | — |
| mistral-small-2506 (closed-source) | — | unknown | — |

## Acknowledgments

Enrico Bono is supported by a grant from the European Union’s Horizon Europe Marie Skłodowska-Curie Actions Doctoral Networks program project PICKED (HORIZON–MSCA–2023–DN-01, grant number 101168626). Louiza Galou is supported by a grant from the European Union’s Horizon Europe Marie Skłodowska-Curie Actions Doctoral Networks Industrial Doctorates program project PROMOTE (HORIZON–MSCA–2023–DN-01, grant number 101169245). Paul Perco and Matthias Ley are members of and would like to cordially thank the COST Action PerMediK, CA21165, supported by COST (European Cooperation in Science and Technology). We gratefully acknowledge Dorota Wojenska for her support in optimizing the overview workflow figure (Figure 1).

## Author contributions

Conceptualization: M.L., P.P., E.B. and F.B.; Methodology: M.L., P.P., E.B. and F.B.; Software: M.L.; Formal analysis: F.B., E.B., P.P. and M.L.; Validation: E.B., F.B., L.F., P.P., M.L., L.G., K.K.-I., S.W., P.A. and K.K.; Writing—original draft: F.B., E.B., M.L. and P.P.; Writing—review & editing: E.B., F.B., P.P., M.L., K.K., L.G., L.F., K.K.-I., P.A. and S.W.; Visualization: F.B. and E.B.; Supervision: M.L. and P.P.; All authors have read and agreed to the published version of the manuscript.

## Declaration of generative AI and AI-assisted technologies in the writing process

During the preparation of this work the author(s) used Anthropic (Claude Opus 4.5) and OpenAI (GPT-5) in order to enhance textual clarity and readability without introducing of new hypotheses or data. After using this tool/service, the author(s) reviewed and edited the content as needed and take(s) full responsibility for the content of the published article.

## References

1. Zhang, Y., Jin, H., Meng, D., Wang, J., and Tan, J. (2025). A comprehensive survey on process-oriented automatic text summarization with exploration of llm-based methods. URL: https://arxiv.org/abs/2403.02901. arXiv:2403.02901.

2. Zhang, H., Yu, P. S., and Zhang, J. (2024). A systematic survey of text summarization: From statistical methods to large language models. URL: https://arxiv.org/abs/2406.11289. arXiv:2406.11289.

3. Rohil, M. K., and Magotra, V. (2022). An exploratory study of automatic text summarization in biomedical and healthcare domain. Healthcare Analytics 2, 100058. URL: https://www.sciencedirect.com/science/article/pii/S2772442522000223. 10.1016/j.health.2022.100058.

4. Xie, Q., Luo, Z., Wang, B., and Ananiadou, S. (2023). A survey for biomedical text summarization: From pre-trained to large language models. URL: https://arxiv.org/abs/2304.08763. arXiv:2304.08763.

5. Chen, Q., Hu, Y., Peng, X., Xie, Q., Jin, Q., Gilson, A., Singer, M. B., Ai, X., Lai, P.-T., Wang, Z., Keloth, V. K., Raja, K., Huang, J., He, H., Lin, F., Du, J., Zhang, R., Zheng, W. J., Adelman, R. A., Lu, Z., and Xu, H. (2025). Benchmarking large language models for biomedical natural language processing applications and recommendations. Nature Communications 16, 3280. URL: 10.1038/s41467-025-56989-2. doi:10.1038/s41467-025-56989-2.

6. Celikten, T., and Onan, A. (2025). Benchmarking large language models for biomedical literature summarization: Abstractive versus extractive paradigms. IEEE Access 13, 152682–152715. URL: 10.1109/ACCESS.2025.3604351. doi:10.1109/ ACCESS.2025.3604351.

7. Toprak, A. G., and Onan, A. (2025). Benchmarking intelligent large language models for biomedical text summarization: A performance evaluation. In: Intelligent and Fuzzy Systems (INFUS 2025) vol. 1529 of Lecture Notes in Networks and Systems. Springer (492–500). doi:10.1007/978-3-031-97992-7_55.

8. Luhn, H. P. (1958). The automatic creation of literature abstracts. IBM J. Res. Dev. 2, 159–165. URL: https://api.semanticscholar.org/CorpusID:15475171.

9. Edmundson, H. P. (1969). New methods in automatic extracting. J. ACM 16, 264–285. URL: 10.1145/321510.321519. doi:10.1145/321510.321519.

10. Robertson, S. (2004). Understanding inverse document frequency: On theoretical arguments for idf. Journal of Documentation - J DOC 60, 503–520. doi:10.1108/00220410410560582.

11. Reeve, L. H., Han, H., Nagori, S. V., Yang, J. C., Schwimmer, T. A., and Brooks, A. D. (2006). Concept frequency distribution in biomedical text summarization. In: Proceedings of the 15th ACM International Conference on Information and Knowledge Management. CIKM ’06 New York, NY, USA: Association for Computing Machinery. ISBN 1595934332 (604–611). URL: 10.1145/1183614.1183701. doi:10.1145/1183614.1183701.

12. Mihalcea, R., and Tarau, P. (2004). TextRank: Bringing order into text. In: Lin, D., and Wu, D., eds. Proceedings of the 2004 Conference on Empirical Methods in Natural Language Processing. Barcelona, Spain: Association for Computational Linguistics (404–411). URL: https://aclanthology.org/W04-3252/.

13. Afzal, M., Alam, F., Malik, K. M., and Malik, G. M. (2020). Clinical context–aware biomedical text summarization using deep neural network: Model development and validation. J Med Internet Res 22, e19810. URL: http://www.jmir.org/2020/10/e19810/. doi:10.2196/19810.

14. Almasoud, A., Hassine, S., Al-Wesabi, F., Nour, M., Hilal, A., Al Duhayyim, M., Hamza, A., and Motwakel, A. (2022). Automated multi-document biomedical text summarization using deep learning model. Computers, Materials & Continua 71, 5800. doi:10.32604/cmc.2022.024556.

15. Vaswani, A., Shazeer, N., Parmar, N., Uszkoreit, J., Jones, L., Gomez, A. N., Kaiser, L., and Polosukhin, I. (2023). Attention is all you need. URL: https://arxiv.org/abs/1706.03762. arXiv:1706.03762.

16. Devlin, J., Chang, M.-W., Lee, K., and Toutanova, K. (2019). Bert: Pre-training of deep bidirectional transformers for language understanding. URL: https://arxiv.org/abs/1810.04805. arXiv:1810.04805.

17. Lewis, M., Liu, Y., Goyal, N., Ghazvininejad, M., Mohamed, A., Levy, O., Stoyanov, V., and Zettlemoyer, L. (2019). BART: denoising sequence-to-sequence pre-training for natural language generation, translation, and comprehension. CoRR abs/1910.13461. URL: http://arxiv.org/abs/1910.13461. arXiv:1910.13461.

18. Yuan, H., Yuan, Z., Gan, R., Zhang, J., Xie, Y., and Yu, S. (2022). BioBART: Pretraining and evaluation of a biomedical generative language model. In: Demner-Fushman, D., Cohen, K. B., Ananiadou, S., and Tsujii, J., eds. Proceedings of the 21st Workshop on Biomedical Language Processing. Dublin, Ireland: Association for Computational Linguistics (97–109). URL: https://aclanthology.org/2022.bionlp-1.9/. doi:10.18653/v1/2022.bionlp-1.9.

19. Abinaya, S., Vigil, M., Keerthika, K., and Varshasri, R. (2024). Medical text summarization using bart with lora-based parameter efficient fine tuning.

20. Raffel, C., Shazeer, N., Roberts, A., Lee, K., Narang, S., Matena, M., Zhou, Y., Li, W., and Liu, P. J. (2020). Exploring the limits of transfer learning with a unified text-to-text transformer. Journal of Machine Learning Research 21, 1–67. URL: http://jmlr.org/papers/v21/20-074.html.

21. Zhang, J., Zhao, Y., Saleh, M., and Liu, P. J. (2020). Pegasus: Pre-training with extracted gap-sentences for abstractive summarization. URL: https://arxiv.org/abs/1912.08777. arXiv:1912.08777.

22. Beltagy, I., Peters, M. E., and Cohan, A. (2020). Longformer: The long-document transformer. URL: https://arxiv.org/abs/2004.05150. arXiv:2004.05150.

23. Steblianko, O., Shymkovych, V., Kravets, P., Novatskyi, A., and Shymkovych, L. (2024). Scientific article summarization model with unbounded input length. Information, Computing and Intelligent systems (150–158). doi:10.20535/2786-8729.5.2024.314724.

24. Radford, A., and Narasimhan, K. (2018). Improving language understanding by generative pre-training. URL: https://api.semanticscholar.org/CorpusID:49313245.

25. Bai, Y., Kadavath, S., Kundu, S., Askell, A., Kernion, J., Jones, A., Chen, A., Goldie, A., Mirhoseini, A., McKinnon, C., et al. (2022). Constitutional ai: Harmlessness from ai feedback. URL: https://arxiv.org/abs/2212.08073. arXiv:2212.08073.

26. Grattafiori, A., Dubey, A., Jauhri, A., Pandey, A., Kadian, A., Al-Dahle, A., Letman, A., Mathur, A., Schelten, A., Vaughan, A., et al. (2024). The llama 3 herd of models. URL: https://arxiv.org/abs/2407.21783. arXiv:2407.21783.

27. Team, G., Kamath, A., Ferret, J., Pathak, S., Vieillard, N., Merhej, R., Perrin, S., Mate-jovicova, T., Ramé, A., Rivière, M., et al. (2025). Gemma 3 technical report. URL: https://arxiv.org/abs/2503.19786. arXiv:2503.19786.

28. Plaat, A., Wong, A., Verberne, S., Broekens, J., van Stein, N., and Back, T. (2025). Multi-step reasoning with large language models, a survey. URL: https://arxiv.org/abs/2407.11511. arXiv:2407.11511.

29. Wang, C., and Kantarcioglu, M. (2025). A review of deepseek models’ key innovative techniques. URL: https://arxiv.org/abs/2503.11486. arXiv:2503.11486.

30. Team, Q. (2025). Qwen3 technical report. URL: https://arxiv.org/abs/2505.09388. arXiv:2505.09388.

31. Mistral-AI, :, Rastogi, A., Jiang, A. Q., Lo, A., Berrada, G., Lample, G., Rute, J., Barmentlo, J., Yadav, K., Khandelwal, K., et al. (2025). Magistral. URL: https://arxiv.org/abs/2506.10910. arXiv:2506.10910.

32. Ankit Pal, M. S. (2024). Openbiollms: Advancing open-source large language models for healthcare and life sciences. https://huggingface.co/aaditya/OpenBioLLM-Llama3-70B Hugging Face.

33. Luo, R., Sun, L., Xia, Y., Qin, T., Zhang, S., Poon, H., and Liu, T.-Y. (2022). BioGPT: generative pre-trained transformer for biomedical text generation and mining. Briefings in Bioinformatics 23. URL: 10.1093/bib/bbac409. doi:10.1093/bib/bbac409. arXiv:https://academic.oup.com/bib/article-pdf/23/6/bbac409/47144271/bbac409.pdf.Bbac409.

34. Labrak, Y., Bazoge, A., Morin, E., Gourraud, P.-A., Rouvier, M., and Dufour, R. (2024). Biomistral: A collection of open-source pretrained large language models for medical domains. arXiv:2402.10373.

35. Li, S., Huang, J., Zhuang, J., Shi, Y., Cai, X., Xu, M., Wang, X., Zhang, L., Ke, G., and Cai, H. (2025). Scilitllm: How to adapt llms for scientific literature understanding. URL: https://arxiv.org/abs/2408.15545. arXiv:2408.15545.

36. Gallifant, J., Afshar, M., Ameen, S., Aphinyanaphongs, Y., Chen, S., Cacciamani, G., Demner-Fushman, D., Dligach, D., Daneshjou, R., Fernandes, C., Hansen, L., Landman, A., Lehmann, L., McCoy, L., Miller, T., Moreno, A., Munch, N., Restrepo, D., Savova, G., Umeton, R., Gichoya, J., Collins, G., Moons, K., Celi, L., and Bitterman, D. (2025). The tripod-llm reporting guideline for studies using large language models. Nature medicine 31, 60–69. doi:10.1038/s41591-024-03425-5. Publisher Copyright: © The Author(s), under exclusive licence to Springer Nature America, Inc. 2025.

37. Elsevier (2024). Highlights. https://www.elsevier.com/researcher/author/tools-and-resources/highlights (accessed: 2025-08-07).

38. Cell Press (2024). Final submission: Other components: Highlights. https://www.cell.com/cell/information-for-authors/final-submission (accessed: 2025-08-07).

39. Belcak, P., Heinrich, G., Diao, S., Fu, Y., Dong, X., Muralidharan, S., Lin, Y. C., and Molchanov, P. (2025). Small language models are the future of agentic ai. URL: https://arxiv.org/abs/2506.02153. arXiv:2506.02153.

40. Lin, C.-Y. (2004). ROUGE: A package for automatic evaluation of summaries. In: Text Summarization Branches Out. Barcelona, Spain: Association for Computational Linguistics (74–81). URL: https://aclanthology.org/W04-1013/.

41. Papineni, K., Roukos, S., Ward, T., and Zhu, W.-J. (2002). Bleu: a method for automatic evaluation of machine translation. In: Isabelle, P., Charniak, E., and Lin, D., eds. Proceedings of the 40th Annual Meeting of the Association for Computational Linguistics. Philadelphia, Pennsylvania, USA: Association for Computational Linguistics (311–318). URL: https://aclanthology.org/P02-1040/. doi:10.3115/1073083.1073135.

42. Banerjee, S., and Lavie, A. (2005). METEOR: An automatic metric for MT evaluation with improved correlation with human judgments. In: Goldstein, J., Lavie, A., Lin, C.-Y., and Voss, C., eds. Proceedings of the ACL Workshop on Intrinsic and Extrinsic Evaluation Measures for Machine Translation and/or Summarization. Ann Arbor, Michigan: Association for Computational Linguistics (65–72). URL: https://aclanthology.org/W05-0909/.

43. Zhang, T., Kishore, V., Wu, F., Weinberger, K. Q., and Artzi, Y. (2020). Bertscore: Evaluating text generation with bert. URL: https://arxiv.org/abs/1904.09675. arXiv:1904.09675.

44. Liu, Y., Ott, M., Goyal, N., Du, J., Joshi, M., Chen, D., Levy, O., Lewis, M., Zettlemoyer, L., and Stoyanov, V. (2019). Roberta: A robustly optimized bert pretraining approach. URL: https://arxiv.org/abs/1907.11692. arXiv:1907.11692.

45. He, P., Liu, X., Gao, J., and Chen, W. (2021). Deberta: Decoding-enhanced bert with disentangled attention. URL: https://arxiv.org/abs/2006.03654. arXiv:2006.03654.

46. Song, K., Tan, X., Qin, T., Lu, J., and Liu, T.-Y. (2020). Mpnet: Masked and permuted pre-training for language understanding. URL: https://arxiv.org/abs/2004. 09297. arXiv:2004.09297.

47. Zha, Y., Yang, Y., Li, R., and Hu, Z. (2023). Alignscore: Evaluating factual consistency with a unified alignment function. URL: https://arxiv.org/abs/2305.16739. arXiv:2305.16739.

48. Laban, P., Schnabel, T., Bennett, P. N., and Hearst, M. A. (2021). Summac: Re-visiting nli-based models for inconsistency detection in summarization. URL: https://arxiv.org/abs/2111.09525. arXiv:2111.09525.

49. Tang, L., Laban, P., and Durrett, G. (2024). Minicheck: Efficient fact-checking of llms on grounding documents. URL: https://arxiv.org/abs/2404.10774. arXiv:2404.10774.

50. GitHub (2026). Llm knowledge cut-off dates summary. https://github.com/HaoooWang/llm-knowledge-cutoff-dates (accessed: 2026-04-09).

51. Fabbri, A. R., Krysćiñski, W., McCann, B., Xiong, C., Socher, R., and Radev, D. (2021). Summeval: Re-evaluating summarization evaluation. URL: https://arxiv.org/abs/2007.12626. arXiv:2007.12626.

52. Liu, Y., Iter, D., Xu, Y., Wang, S., Xu, R., and Zhu, C. (2023). G-eval: Nlg evaluation using gpt-4 with better human alignment. URL: https://arxiv.org/abs/2303.16634. arXiv:2303.16634.

53. Gwet, K. Handbook of inter-rater reliability: The definitive guide to measuring the extent of agreement among raters (2012).

54. Zheng, L., Chiang, W.-L., Sheng, Y., Zhuang, S., Wu, Z., Zhuang, Y., Lin, Z., Li, Z., Li, D., Xing, E., Zhang, H., Gonzalez, J., and Stoica, I. (2023). Judging LLM-as-a-judge with MT-bench and chatbot arena. In: Oh, A., Naumann, T., Globerson, A., Saenko, K., Hardt, M., and Levine, S., eds. Advances in Neural Information Processing Systems vol. 36. Curran Associates, Inc. (46595–46623).

55. Verga, P., Hofstatter, S., Althammer, S., Su, Y., Piktus, A., Arkhangorodsky, A., Xu, M., White, N., and Lewis, P. (2024). Replacing Judges with Juries: Evaluating LLM Generations with a Panel of Diverse Models. arXiv. doi:10.48550/arXiv.2404.18796. arXiv:2404.18796.

56. Liu, Y., Iter, D., Xu, Y., Wang, S., Xu, R., and Zhu, C. (2023). G-Eval: NLG Evaluation using Gpt-4 with Better Human Alignment. In: Bouamor, H., Pino, J., and Bali, K., eds. Proceedings of the 2023 Conference on Empirical Methods in Natural Language Processing. Singapore: Association for Computational Linguistics (2511–2522). doi:10.18653/v1/2023.emnlp-main.153.

57. McKinney, W. (2010). Data structures for statistical computing in python. In: van der Walt, S., and Millman, J., eds. Proceedings of the 9th Python in Science Conference. (51 – 56).

58. Pedregosa, F., Varoquaux, G., Gramfort, A., Michel, V., Thirion, B., Grisel, O., Blondel, M., Prettenhofer, P., Weiss, R., Dubourg, V. et al. (2011). Scikit-learn: Machine learning in Python. Journal of Machine Learning Research 12, 2825–2830.

59. Hagberg, A., Swart, P., and S Chult, D. Exploring network structure, dynamics, and function using networkx. Tech. Rep. Los Alamos National Lab.(LANL), Los Alamos, NM (United States) (2008).

60. Brin, S., and Page, L. (1998). The anatomy of a large-scale hypertextual web search engine. Computer Networks and ISDN Systems 30, 107–117. URL: https://www.sciencedirect.com/science/article/pii/S016975529800110X. doi:10.1016/S0169-7552(98)00110-X. Proceedings of the Seventh International World Wide Web Conference.

61. Bird, S., Klein, E., and Loper, E. Natural language processing with Python: analyzing text with the natural language toolkit. ” O’Reilly Media, Inc.” (2009).

62. Reimers, N., and Gurevych, I. (2019). Sentence-bert: Sentence embeddings using siamese bert-networks. URL: https://arxiv.org/abs/1908.10084. arXiv:1908.10084.

63. Wolf, T., Debut, L., Sanh, V., Chaumond, J., Delangue, C., Moi, A., Cistac, P., Rault, T., Louf, R., Funtowicz, M., et al. (2020). Transformers: State-of-the-art natural language processing. In: Proceedings of the 2020 Conference on Empirical Methods in Natural Language Processing: System Demonstrations. Online: Association for Computational Linguistics (38–45). URL: https://www.aclweb.org/anthology/2020.emnlp-demos.6.

64. Panickssery, A., Bowman, S. R., and Feng, S. (2024). LLM Evaluators Recognize and Favor Their Own Generations. In: Advances in Neural Information Processing Systems vol. 37. Curran Associates, Inc. (68772–68802). doi:10.52202/079017-2197.

65. Pombal, J., Rei, R., and Martins, A. F. T. (2026). Self-Preference Bias in Rubric-Based Evaluation of Large Language Models. arXiv. doi:10.48550/arXiv.2604.06996. arXiv:2604.06996.

66. Xu, B., Chen, Y., Wen, Z., Liu, W., and He, B. (2025). Evaluating small language models for news summarization: Implications and factors influencing performance. URL: https://arxiv.org/abs/2502.00641. arXiv:2502.00641.

67. Muennighoff, N., Rush, A. M., Barak, B., Scao, T. L., Piktus, A., Tazi, N., Pyysalo, S., Wolf, T., and Raffel, C. (2025). Scaling data-constrained language models. URL: https://arxiv.org/abs/2305.16264. arXiv:2305.16264.

68. Dorfner, F. J., Dada, A., Busch, F., Makowski, M. R., Han, T., Truhn, D., Kleesiek, J., Sushil, M., Lammert, J., Adams, L. C., and Bressem, K. K. (2024). Biomedical large languages models seem not to be superior to generalist models on unseen medical data. URL: https://arxiv.org/abs/2408.13833. arXiv:2408.13833.

69. Zhou, Y., Liu, X., Zhang, X., Ning, C., Wang, S., Hu, G., and Wu, J. (2025). Investigating and mitigating catastrophic forgetting in medical knowledge injection through internal knowledge augmentation learning. OpenReview. Preprint. Available online: https://openreview.net/forum?id=i9RDDi2SZC.

70. Jin, K., Wang, Y., Santos, L., Fang, T., Yang, X., Im, S. K., and Oliveira, H. G. (2025). Reasoning or not? a comprehensive evaluation of reasoning llms for dialogue summarization. URL: https://arxiv.org/abs/2507.02145. arXiv:2507.02145.

71. Durmus, E., He, H., and Diab, M. (2020). FEQA: A question answering evaluation framework for faithfulness assessment in abstractive summarization. In: Jurafsky, D., Chai, J., Schluter, N., and Tetreault, J., eds. Proceedings of the 58th Annual Meeting of the Association for Computational Linguistics. Online: Association for Computational Linguistics (5055–5070). URL: https://aclanthology.org/2020.acl-main.454/. doi:10.18653/v1/2020.acl-main.454.

72. Irugalbandara, C., Mahendra, A., Daynauth, R., Arachchige, T. K., Dantanarayana, J., Flautner, K., Tang, L., Kang, Y., and Mars, J. (2024). Scaling down to scale up: A cost-benefit analysis of replacing openai’s llm with open source slms in production. URL: https://arxiv.org/abs/2312.14972. arXiv:2312.14972.

73. Khattab, O., Singhvi, A., Maheshwari, P., Zhang, Z., Santhanam, K., Vardhamanan, S., Haq, S., Sharma, A., Joshi, T. T., Moazam, H., Miller, H., Zaharia, M., and Potts, C. (2023). Dspy: Compiling declarative language model calls into self-improving pipelines. URL: https://arxiv.org/abs/2310.03714.arXiv:2310.03714.

74. Khattab, O., Santhanam, K., Li, X. L., Hall, D., Liang, P., Potts, C., and Zaharia, M. (2023). Demonstrate-search-predict: Composing retrieval and language models for knowledge-intensive nlp. URL: https://arxiv.org/abs/2212.14024. arXiv:2212.14024.

75. Kohli, G. (2026). Nine Judges, Two Effective Votes: Correlated Errors Undermine LLM Evaluation Panels. arXiv. doi:10.48550/arXiv.2605.29800. arXiv:2605.29800.

